# Human-specific NOTCH2NL promotes astrogenesis by expanding proliferative glial progenitor states

**DOI:** 10.64898/2025.12.16.694764

**Authors:** Riina Ishiwatari, Xuanhao D. Sheu, Rintaro Amano, Yuki Y. Yamauchi, Pauline Rouillard, Takuma Kumamoto, Yusuke Kishi, Kazuo Emoto, Ikuo K. Suzuki

## Abstract

The human cerebral cortex contains an unusually large number of glial cells, particularly astrocytes, yet the developmental and genetic mechanisms underlying their expansion remain poorly understood. While human-specific genes have been shown to promote neuronal production during cortical development, whether such genes also regulate gliogenesis has remained unclear. Here, we identify a previously unrecognized role for the human-specific gene family NOTCH2NL in promoting astrocyte-lineage expansion. Reanalysis of human fetal single-cell transcriptomic datasets revealed that NOTCH2NL is robustly expressed along the gliogenic trajectory, from glial intermediate progenitor cells to astrocytes. Functional perturbations in a human astrocyte culture system demonstrated that NOTCH2NL is both required and sufficient for astrocyte proliferation. In vivo overexpression of NOTCH2NLB in the developing mouse cortex shifted progenitor output toward the astrocyte lineage, increasing the astrocyte-to-neuron ratio from the early postnatal period through adulthood. This phenotype was associated with an expansion of proliferative glial progenitors around birth. Single-nucleus transcriptomic profiling further showed that NOTCH2NLB suppresses neuronal gene programs while activating transcriptional modules related to cell proliferation and cellular homeostasis during gliogenesis. Together, these findings indicate that human-specific NOTCH2NL acts at a conserved developmental decision point to amplify astrocyte production. Our study extends the function of NOTCH2NL beyond neurogenesis and suggests that human lineage–specific gene duplications can modulate gliogenesis, providing a developmental mechanism that may have contributed to the coordinated expansion of neuronal and glial populations in the human cortex.

## Introduction

The generation of glial cells is a fundamental step in cortical development, essential for circuit formation, metabolic support, and long-term brain function. In contrast to neurogenesis, whose cellular and molecular regulation has been extensively characterized, the mechanisms that control the timing, magnitude, and lineage commitment of gliogenesis remain comparatively poorly understood. In particular, how radial glial (RG) progenitors transition from neurogenic programs to gliogenic programs and how this transition is quantitatively regulated remain a central open question in developmental neurobiology.

Astrocytes constitute a major glial population in the mammalian cortex and play critical roles in synaptic regulation, ion homeostasis, and neuronal metabolism <u>(Khakh and Deneen, 2019; Oberheim et al., 2006; Sloan and Barres, 2014; Zuchero and Barres, 2015)</u>. Their numbers are tightly regulated during development, yet the mechanisms that determine astrocyte output are incompletely defined. In rodents, astrocytes are generated after neurogenesis through intermediate progenitor states, including glial intermediate progenitor cells (glial-IPCs, also termed tripotent IPCs), which amplify astrocyte production through proliferative divisions. Although several signaling pathways, including Notch signaling, have been implicated in astrocyte specification, the specific progenitor stages at which these pathways act, and how progenitor proliferation versus lineage commitment is coordinated during gliogenesis, remain unresolved.

These questions are particularly salient in the context of the human cerebral cortex, where astrocytes are more abundant and exhibit distinct morphological and functional properties compared to those in rodents. The glia-to-neuron ratio is substantially higher in humans than in many other mammals, suggesting that quantitative regulation of gliogenesis is a key variable in cortical development <u>(Herculano-Houzel, 2014; Oberheim Bush and Nedergaard, 2017)</u>. Notably, astrocytes display some of the most pronounced species-specific features, including increased size, enhanced process complexity, and extensive overlap between neighboring cells <u>(Oberheim et al., 2009, 2006)</u>. Despite their importance, the molecular mechanisms that drive increased astrocyte production, both in general and in a species-dependent manner, remain largely unknown.

Recent work has highlighted that human lineage–specific gene duplications can modulate core developmental programs. One prominent example is NOTCH2NL (N2NL), a family of paralogs generated by partial duplication of the highly conserved NOTCH2 receptor gene after divergence from chimpanzees <u>(Fiddes et al., 2018; Florio et al., 2018; Suzuki et al., 2018)</u>. N2NL enhances Notch signaling and prolongs the maintenance of neural progenitors during cortical neurogenesis, thereby increasing neuronal output and cortical size. These findings establish N2NL as a regulator of progenitor dynamics during neurogenesis. However, whether this human-specific gene also regulates gliogenic programs and, if so, at which progenitor stages, has remained unexplored.

Glial cells are produced following neurons during cortical development. In mice, embryonic ventricular radial glia (vRG) generate excitatory neurons either directly or via neuronal intermediate progenitors. After neurogenesis, a subset of vRG transitions into glial-IPCs, which express markers such as EGFR and Olig2 and give rise to astrocytes, oligodendrocytes, and inhibitory neurons <u>(Li et al., 2021; Zhang et al., 2020)</u>. In the human cortex, gliogenesis proceeds through both conserved and divergent routes, involving truncated RG (tRG), outer RG (oRG), and EGFR-positive glial-IPCs, which together contribute to astrocyte production <u>(Allen et al., 2022; Wang et al., 2025)</u>. These observations raise the possibility that quantitative regulation of progenitor proliferation and fate choice during the RG-to–glial-IPC transition represents a key control point in cortical gliogenesis.

In this study, we asked whether the human-specific gene N2NL regulates gliogenesis by acting on glial progenitor programs. We first analyzed the transcriptional dynamics of N2NL during human fetal cortical development, focusing on the transition from RG to astrocytes. We then performed gain- and loss-of-function experiments in a human astrocyte culture system, followed by *in vivo* overexpression studies in the developing mouse cortex. Finally, we used single-nucleus transcriptomic approaches to identify molecular programs associated with N2NL activity during gliogenesis. Together, our results uncover a previously underappreciated role for N2NL in promoting astrocyte-lineage proliferation and highlight a genetic mechanism that modulates gliogenesis at a conserved developmental decision point.

## Results

### Endogenous expression of N2NL during human fetal gliogenesis

To investigate the potential role of N2NL in human gliogenesis, we reanalyzed publicly available single-cell RNA-seq datasets of human fetal cortices covering developmental stages from early neurogenesis to gliogenesis (Fig. 1A)<u>(Wang et al., 2025)</u>. Because N2NL comprises four highly similar paralogs, individual sequencing reads are less likely to map uniquely, leading to a significant underestimation of absolute expression levels (Suzuki et al., 2018). However, this technical bias is expected to be consistent across samples and therefore should not affect relative comparisons among cell types. Consistent with previous reports <u>(Fiddes et al., 2018; Suzuki et al., 2018)</u>, the combined expression of the four N2NL paralogs (A, B, C, and R) was readily detected in neuroepithelial cells and vRG during the early neurogenic phase in the first trimester. In contrast, during late neurogenesis and gliogenesis in the second and third trimesters, strong expression was not prominent in neural progenitor subtypes such as oRG and tRG, but was evident in glial-IPC and astrocytes (Fig. 1A). Trajectory analysis indicated that N2NL expression remained high along the lineage progressing from RG → glial-IPC → astrocytes, consistent with sustained involvement across glial differentiation and maturation (Fig. 1B). Unbiased correlation analysis further showed that genes whose expression negatively correlated with N2NL were enriched for neuronal functions (e.g., synapse organization, axonogenesis), whereas genes positively correlated with N2NL were enriched for gliogenic programs (Fig. 1C, Table S1). Together, these results suggest that higher N2NL expression is associated with gliogenic transcriptional states and reduced expression of neuronal gene programs.

**Figure 1.**
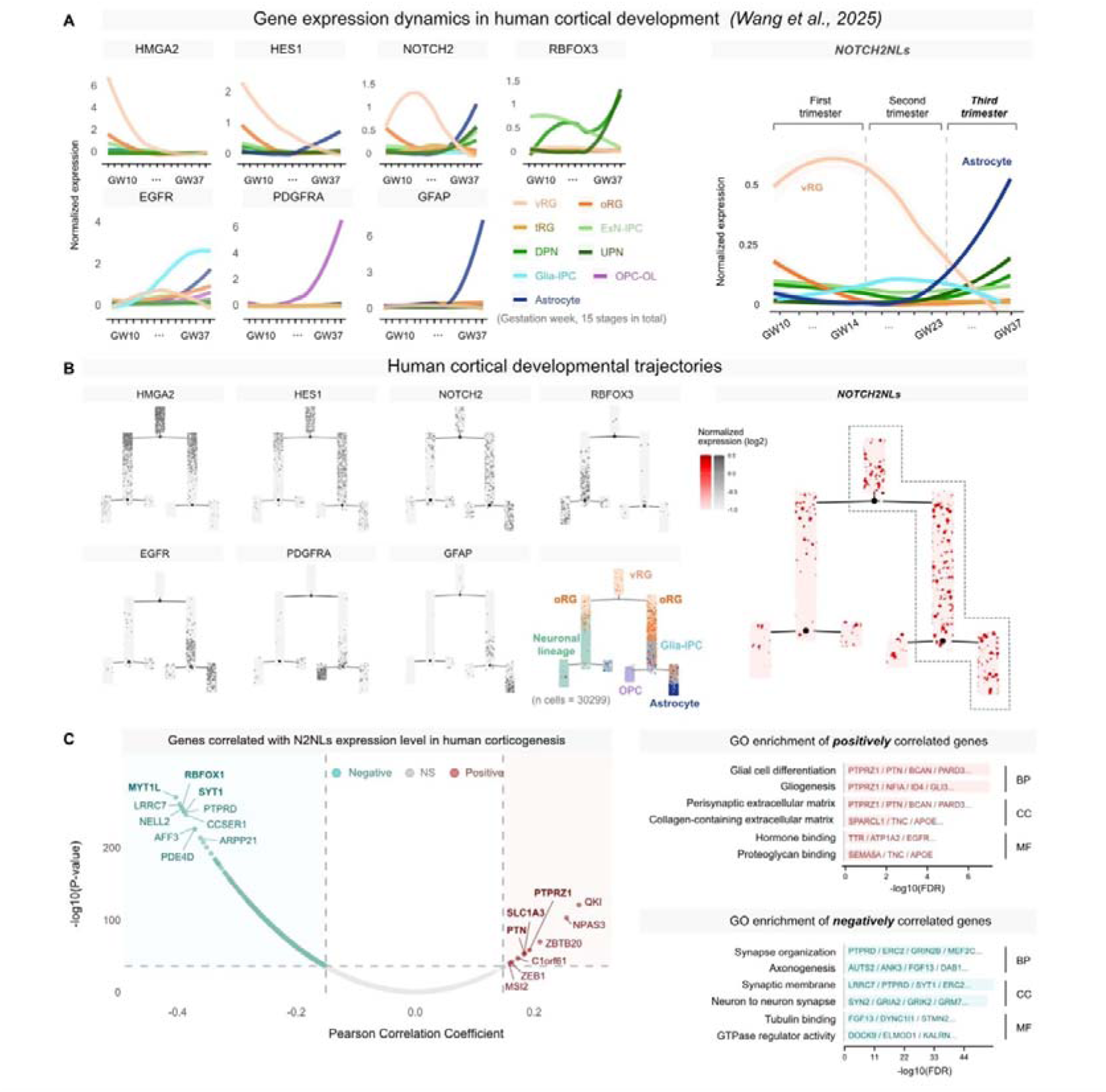
Expression of NOTCH2NL during human fetal corticogenesis. **(A)** Expression dynamics of marker genes for major cortical cell types and the sum of NOTCH2NL paralogs (A, B, C, and R). NOTCH2NL paralogs exhibit strong expression in glial intermediate progenitor cells (glial-IPCs) and astrocytes during the second and third trimesters. **(B)** Monocle2 trajectory analysis reveals a continuous differentiation pathway from vRG to astrocytes. Note the sustained expression of NOTCH2NL along the glial lineage. **(C)** Pearson correlation and Gene Ontology (GO) analysis. The volcano plot (left) shows the correlation of genome-wide gene expression with pooled N2NL levels. Significant correlations (FDR < 0.05, |r| > 0.1) are colored, and the top 10 genes by correlation coefficient are labeled. GO analysis (right) reveals that positively correlated genes are enriched in gliogenic pathways, whereas negatively correlated genes are enriched in neuronal processes. BP, biological process; CC, cellular component; MF, molecular function.

### N2NL promotes the proliferation of human astrocyte culture

We next examined the functional involvement of N2NL in human astrocytes by gain-and loss-of-function analyses. In the culture of immortalized astrocytes derived from human postmortem cortex, lentiviral overexpression of N2NLB, the paralog with the strongest known activity and expression <u>(Fiddes et al., 2018; Suzuki et al., 2018)</u>, significantly increased the proportion of proliferative cells (Ki67^+^ and thymidine analog 5-ethynil-2’-deoxyuridine EdU^+^) at day 5 post-infection (Fig. 2A-C). Conversely, knockdown of N2NLs significantly reduced proliferative populations (Fig. 2D-H). For loss-of-function, short interfering RNA (siRNA) targeting all four N2NL paralogs is introduced into the dissociated human astrocytes with the H2B-mCherry reporter plasmid using 4D-Nucleofector. We validated the efficacy of siRNA by measuring the abundance of N2NL transcripts using RNAscope (Fig. 2D-E). Nucleofected astrocytes were cultured *in vitro* for 4 days, with EdU incorporated during the final 24 hours. The proportion of Ki67- and EdU-positive proliferative cells are quantified in H2B-mCherry-expressing nucleofected cells by immunocytochemistry (Fig. 2F-H). Together, these findings show that N2NL is required for robust proliferation in this astrocyte culture system and that N2NLB overexpression is sufficient to enhance proliferation under these conditions.

**Figure 2.**
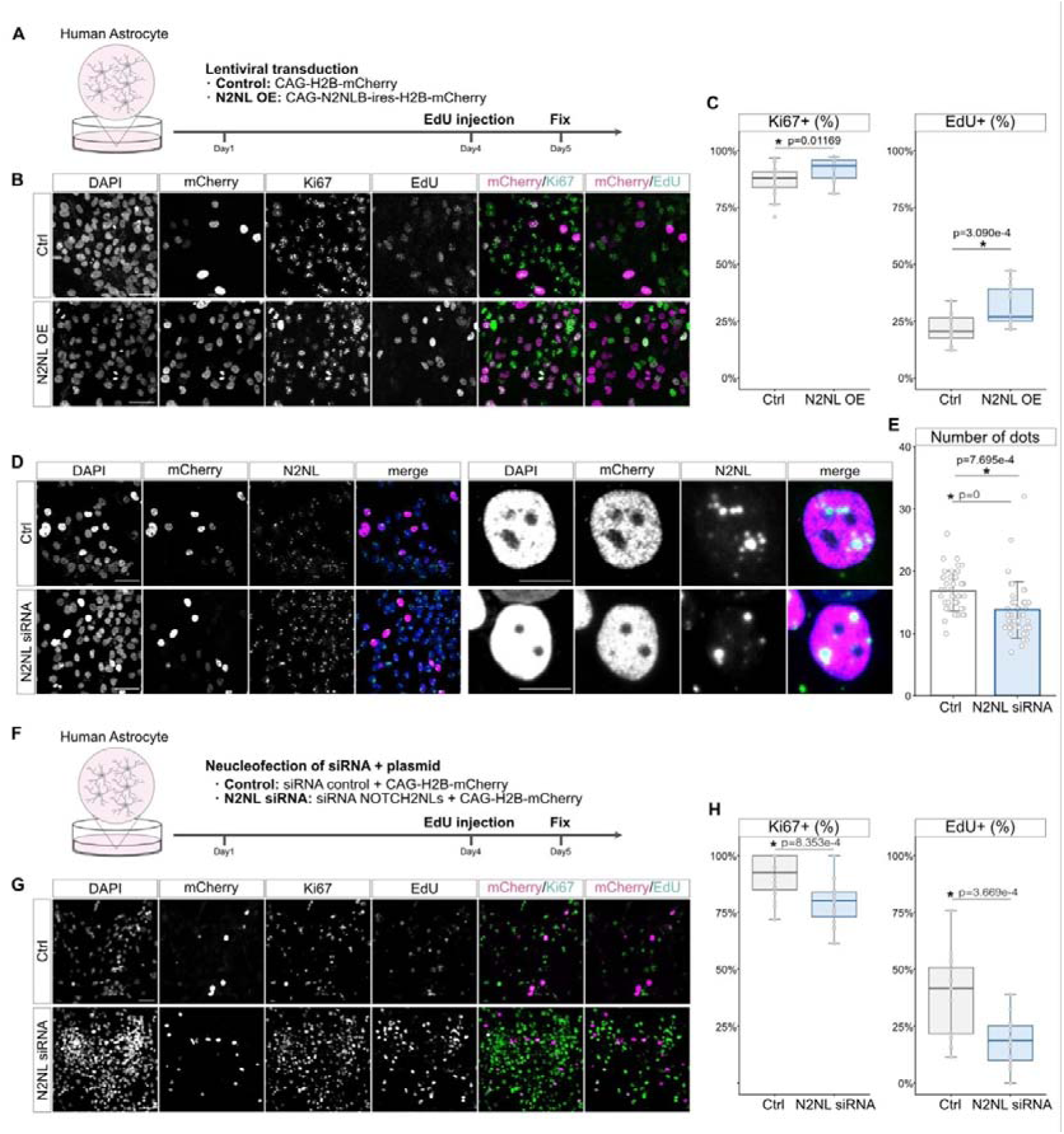
N2NLB regulates the proliferation of human astrocytes. **(A-C)** Lentiviral overexpression of N2NLB promotes astrocyte proliferation. Experimental scheme (**A**), representative immunofluorescence images (**B**), and quantification of Ki67- and EdU-positive cells (**C**) are shown (n = 16 per group). **(D-E)** Validation of knockdown efficiency by RNAscope. Representative images (**D**) and quantification of mRNA dots per cell (**E**) confirm the efficient reduction of *N2NL* mRNA by siRNA (n = 41 cells per group). **(F-H)** siRNA-mediated knockdown of N2NL suppresses astrocyte proliferation. Experimental scheme (**F**), representative images (**G**), and quantification of Ki67- and EdU-positive cells (**H**) are shown (n = 16 wells per group). Statistical analyses: Student’s *t*-test. *, *p* < 0.05. Scale bars: 50 μm.

### N2NLB shifts the neuron-to-astrocyte ratio toward astrocytes in the adult cortex

To investigate the role of N2NL in astrocyte development *in vivo*, we introduced N2NLB into embryonic cortical RGs in mice and assessed astrocytic outcomes in adults. Using *in utero* electroporation at E15.5, we introduced N2NLB into RG progenitors at the ventricular surface of the cortex. Standard plasmid electroporation largely labels postmitotic neurons, as episomal plasmids are diluted by cell division in proliferative cells (Fig. S1A). To enable stable labeling of proliferative cells along the glial differentiation trajectory, we used the iOn-switch system (Kumamoto et al., 2020), which uses a piggyBac transposase to integrate transgenes into the host genome, ensuring persistent expression only after genomic integration (Fig. S1B). To minimize experimental variability, we co-electroporated control mCherry and N2NLB–internal ribosomal entry site (IRES)–EGFP plasmids into the same brains, exploiting the stochastic nature of transposon integration to generate sparse but distinct cell populations within a shared environment (Fig. 3A and Fig. S1B-C). As a proof-of-concept, we confirmed that control mCherry and control EGFP yielded virtually indistinguishable outcomes (Fig. 3B-C).

**Figure 3.**
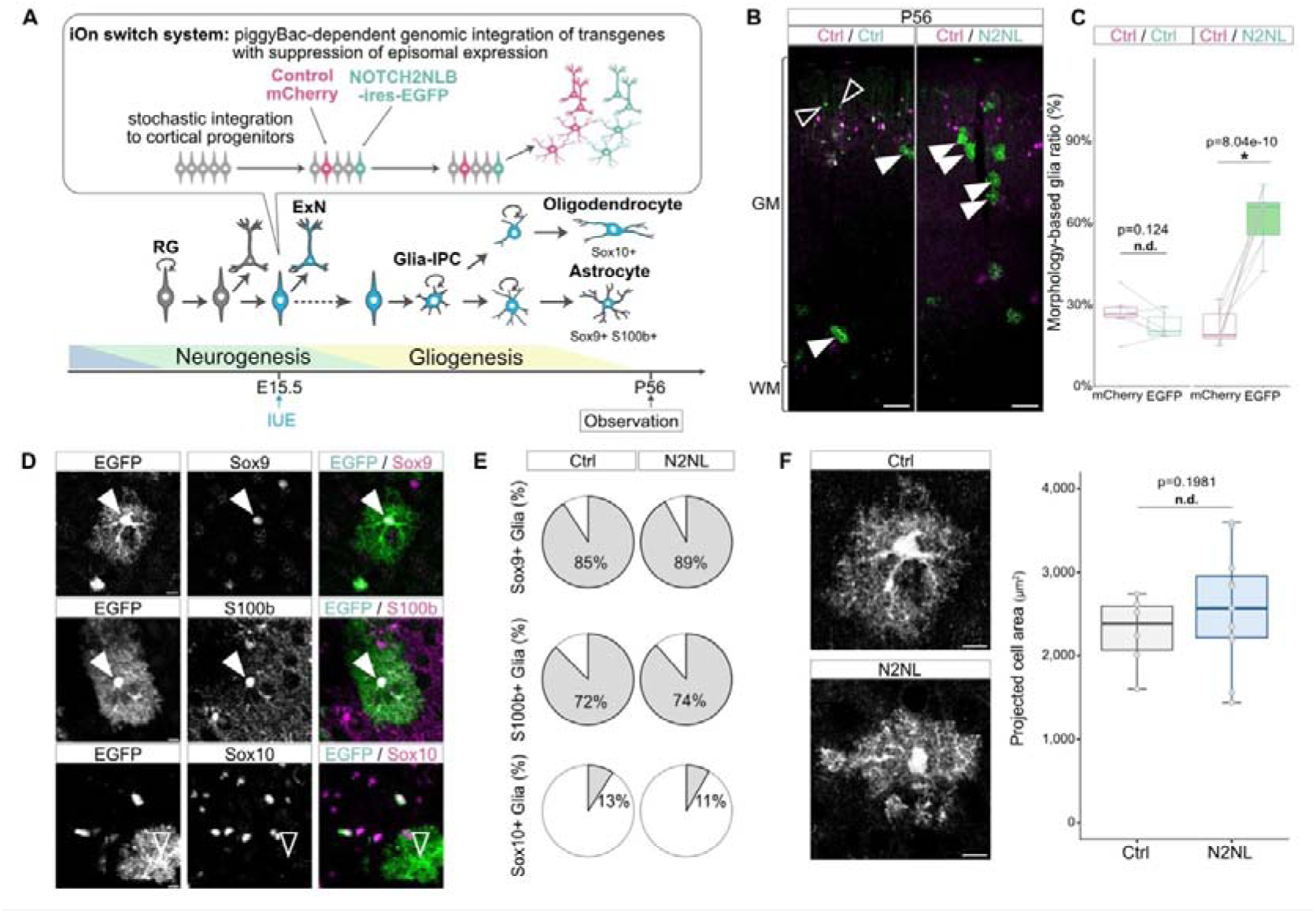
*in vivo* overexpression of N2NLB enhances gliogenesis in mice. **(A)** Schematic representation of the mosaic overexpression experiment using iOn-switch system. Stochastic piggyBac-dependent integration generates mosaic expression of control (mCherry) and N2NLB (EGFP) vectors in cortical progenitors. Cortices were analyzed at P56. **(B-C)** N2NLB increases the proportion of morphologically defined glial cells. Representative images (**B**) and quantification of the glia ratio (**C**) are shown. Filled and open arrowheads indicate morphologically defined glia and neurons, respectively. N2NLB overexpression significantly increases the glial ratio compared to the internal control (Ctrl/Ctrl condition: n = 6 slices; Ctrl/N2NL condition: n = 10 slices). **(D-E)** Characterization of morphologically defined glia with molecular markers. Representative images (**D**) and quantification (**E**) reveal that the majority of morphologically defined glia express astrocytic markers (Sox9, S100b) but are negative for the oligodendrocyte marker Sox10. Filled and open arrowheads indicate marker-positive and -negative cells, respectively. **(F)** Morphological analysis of astrocytes. Representative images (top) and quantification of projected cell area (bottom) show no significant difference in glial size (Ctrl: n = 6 cells; N2NL: n = 11 cells). Statistical analyses: Paired *t*-test (**C**) and Student’s *t*-test (**F**). *, *p* < 0.05; n.d., non-significant difference (*p* ≥ 0.05). Scale bars: 100 μm (**B**) and 10 μm (**D**, **F**).

Overexpression of N2NLB markedly altered the ultimate cell composition of the adult cortex: whereas 23% of mCherry-positive cells exhibited glial morphology in controls, N2NLB increased this fraction to 62% (Fig. 3B-C). Among morphologically identified glial cells in all conditions, ∼90% expressed the astrocyte marker Sox9 and ∼10% expressed the oligodendrocyte marker Sox10, consistent with previous reports indicating that the majority of locally generated cortical glia are astrocytes (Fig. 3D,E)<u>(Clavreul et al., 2019; Shen et al., 2021)</u>. The proportion of astrocytes versus oligodendrocytes within the glial population was not substantially changed by N2NLB overexpression (Fig. 3D-E). We also assessed astrocyte size, since human astrocytes are substantially larger than their murine counterparts. N2NLB-expressing astrocytes showed a small, statistically marginal increase in projected cell area relative to controls (Fig. 3F). These findings indicate that N2NLB enhances astrocyte production *in vivo* without markedly altering gross cell size and glial fate preference.

### N2NLB stimulates the glial progenitor program around birth to amplify astrocyte output

To determine when the astrocyte-lineage bias emerges under N2NLB overexpression, we analyzed cortical tissues during gliogenesis at E18.5, P1, P4, P8, and P15 following E15.5 electroporation (Fig. 4A). Neurons and glia were difficult to distinguish morphologically before P1, but by P4 glial cells could be reliably identified by their characteristic bushy morphology (Fig. 4B and Fig. S2A-B). Across postnatal stages, N2NLB consistently increased the proportion of morphologically defined glial cells relative to controls (Fig. 4B-C), whereas control mCherry and control EGFP showed nearly identical results. More than 80% of morphologically defined glial cells were immunolabeled for the astrocyte marker Sox9 (Fig. 4D), while fewer than 3% of morphologically defined neurons were Sox9-positive (Fig. S2B-C), supporting that morphology-based classification largely corresponds to marker-based identity at P4-P15. These results indicate that the astrocyte-enriched phenotype observed in adults is already established during early postnatal stages.

**Figure 4.**
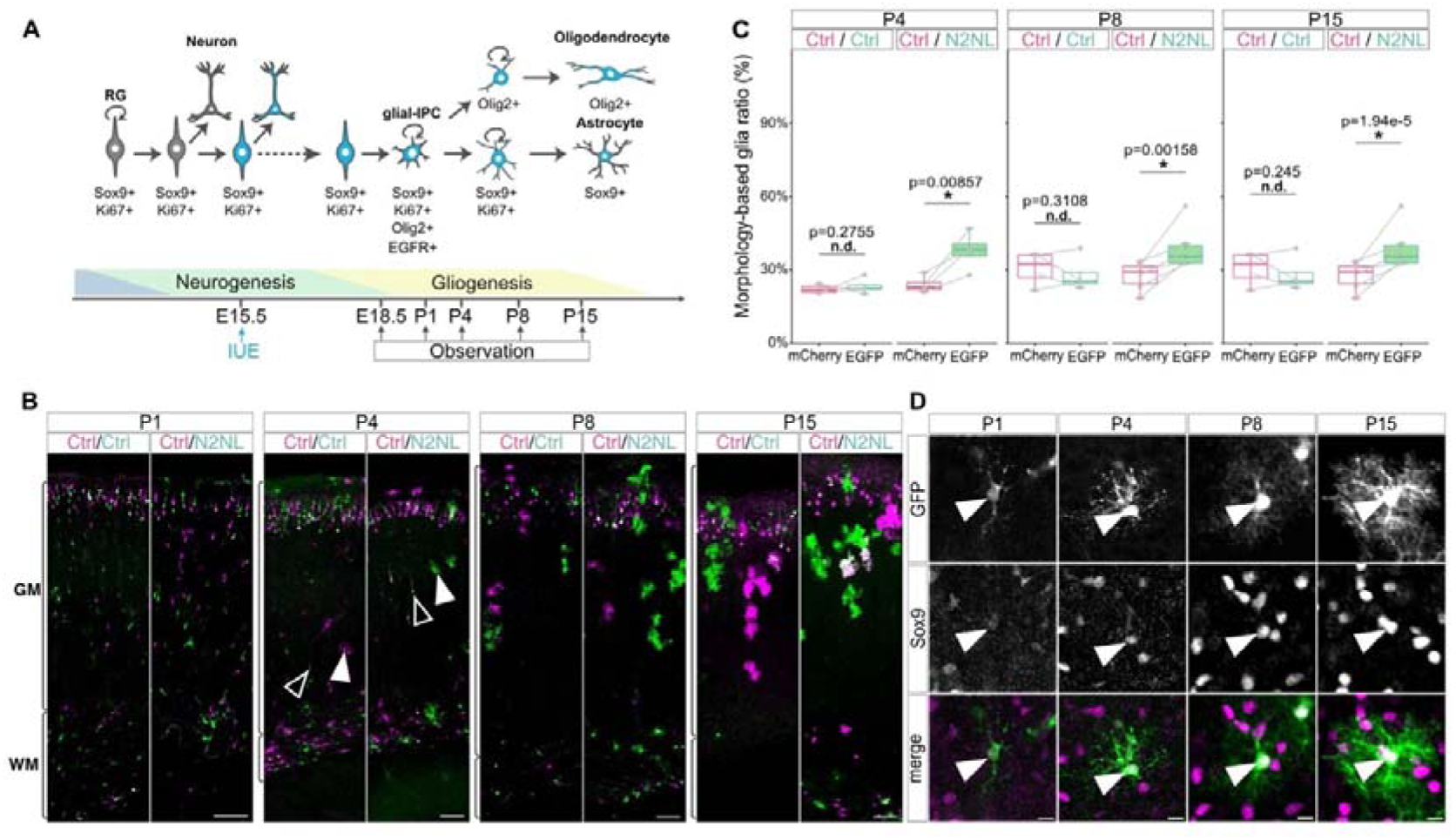
N2NLB increases glial ratio in the early postnatal period. **(A)** Schematic illustration of experimental paradigms. The glial cell ratio among electroporated cells was examined during the gliogenic period in the perinatal and early postnatal stages (P1-P15). **(B)** Representative images of the electroporated cortices at P1, P4, P8, and P15. Glial cells with characteristic bushy morphology are clearly recognizable at P4, P8, and P15 (arrowheads). In contrast, neurons and glia are difficult to distinguish by morphology alone at P1. **(C)** Quantifications of the morphologically defined glia ratio. N2NLB overexpression significantly increases the glial ratio at P4, P8, and P15 compared to the internal control. Note that no significant difference was observed in the Ctrl/Ctrl condition at any stage (P4: Ctrl/Ctrl, n = 4 slices; Ctrl/N2NL, n = 4 slices. P8: Ctrl/Ctrl, n = 4 slices; Ctrl/N2NL, n = 6 slices. P15: Ctrl/Ctrl, n = 10 slices; Ctrl/N2NL, n = 12 slices). **(D)** Validation of cell identity. Representative images showing that the great majority of morphologically defined glial cells are immunolabeled by the astrocytic marker Sox9 (arrowheads). Statistical analyses: Paired *t*-test. *, *p* < 0.05; n.d., non-significant difference (*p* ≥ 0.05). Scale bars: 100 μm (B) and 10 μm (D).

Because astrocytes arise from RG through glial-IPC, which are immunolabeled with Ki67, Olig2, and EGFR (Fig. 5A), we hypothesized that N2NLB expands a proliferative progenitor pool during development to increase astrocyte output in adults. From E18.5 to P8, the proportion of Ki67-positive proliferative cells was consistently increased by N2NLB overexpression (Fig. 5B and Fig. S4). Since Ki67-positive cells include multiple progenitor types across differentiation steps, we classified them into morphologically and molecularly distinct categories. Ki67 cells were detected in both morphologically glial cells and “other” cells (cells that are morphologically neither glial nor neuronal, typically show simpler cell shape with one or two processes, and likely include immature glial precursors, such as glial-IPC and committed immature astrocytes), but were rarely detected in neuronal-shaped cells (Fig. S3). N2NLB did not change the proportion of Ki67 cells within the morphologically glial population, whereas it significantly increased the Ki67 fraction within the “other” population, suggesting a stronger effect on immature glial precursor states than on more mature astrocytes. Olig2 is expressed in glial-IPC and committed oligodendrocyte lineage cells. As most Olig2 cells in these stages also expressed a glial-IPC marker EGFR (Fig. 5A) <u>(Li et al., 2021; Wang et</u> <u>al., 2025)</u>, we used Olig2 as a practical marker to quantify glial-IPC-like populations in this context. N2NLB increased the fraction of Olig2 cells at the onset of cortical gliogenesis at P1 (Fig. 5C). These data support the idea that N2NLB increases adult astrocyte output by expanding proliferative glial progenitors around birth.

**Figure 5.**
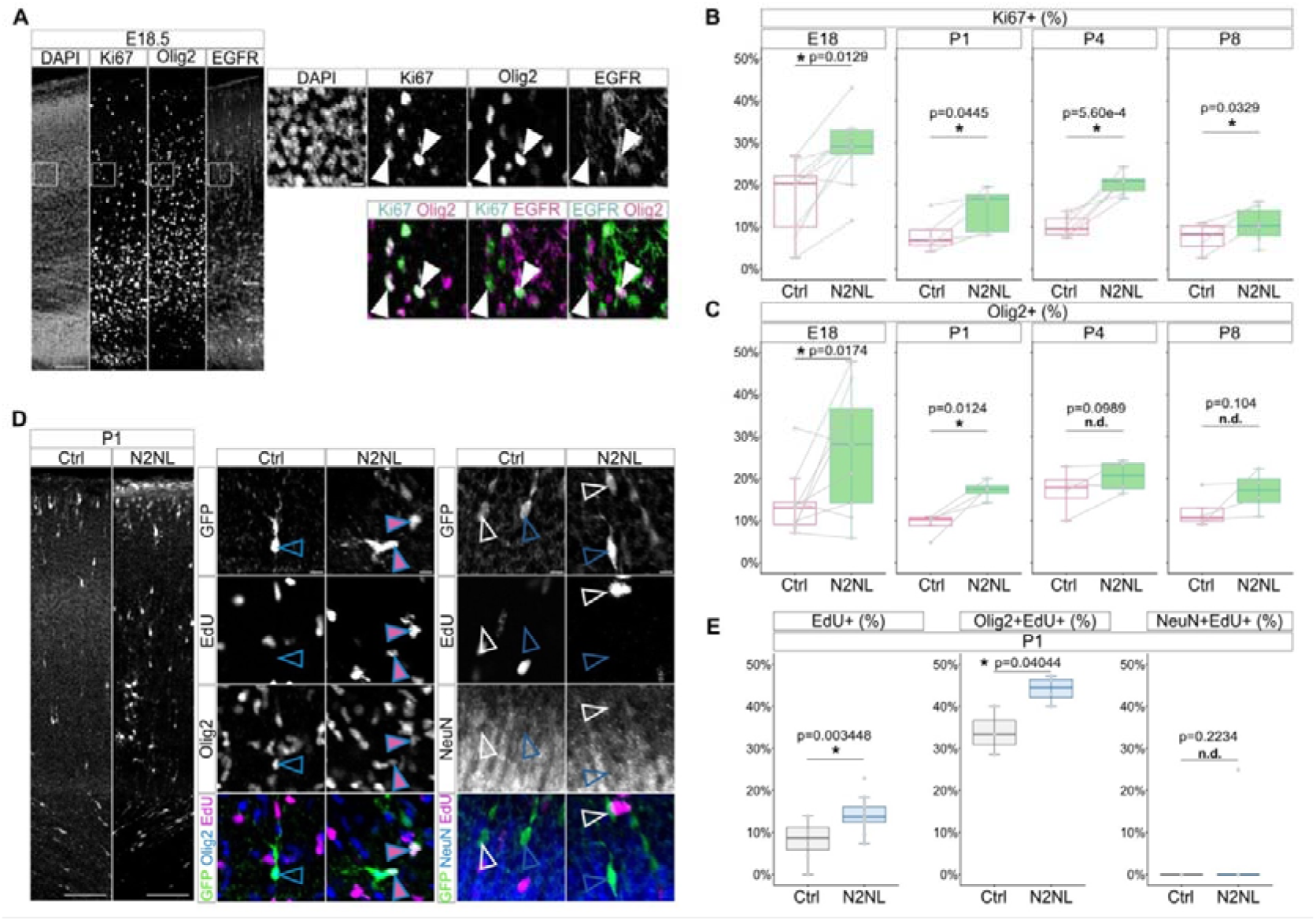
N2NLB induces a higher abundance of glial-IPCs. **(A)** Representative images of glial-IPCs at E18.5, immunolabeled for Ki67, Olig2, and EGFR. Arrowheads indicate triple-positive glial-IPCs. **(B-C)** Quantification of proliferative cells and glial lineage cells. N2NLB overexpression maintains a higher ratio of Ki67-positive proliferative cells (**B**) and Olig2-positive cells (**C**) compared to the control throughout the examined period (Ki67: E18, n = 9 slices; P1, n = 5 slices; P4, n = 6 slices; P8, n = 5 slices. Olig2: E18, n = 9 slices; P1-P8, n = 4 slices). **(D-E)** Fate choice analysis of P0-born cells. (**D**) Representative images of P1 cortices after IUE at E15.5. Blue open arrowheads indicate Olig2+ or NeuN+ electroporated cells. Magenta arrowheads with blue line indicate EdU+ Olig2+ electroporated cells. (**E**) Quantification of EdU incorporation and cell fate among the dividing population. Left: The percentage of EdU-positive cells among the total GFP-positive population is significantly increased in the N2NLB group (Ctrl: n = 8 slices; N2NL: n = 13 slices). Middle & Right: The percentages of Olig2-positive (Middle) and NeuN-positive (Right) cells within the EdU-positive proliferating population (GFP^+^EdU^+^). N2NLB significantly increases the proportion of Olig2-positive glial lineage cells within the dividing population (Ctrl: n = 4 slices; N2NL: n = 7 slices), while the neuronal ratio remains unchanged (Ctrl: n = 4 slices; N2NL: n = 6 slices). Statistical analyses: Paired *t*-test (**B**-**C**) and unpaired Student’s *t*-test (**E**). *, *p* < 0.05; n.d., non-significant difference (*p* ≥ 0.05). Scale bars: 100 μm (**A**, **D** low magnification) and 10 μm (**A**, **D** high magnification).

Previously, we reported that introducing N2NLB into E15.5 vRG progenitors increases the number of Pax6 undifferentiated vRG and decreases the number of βIII tubulin postmitotic neurons at E18.5 <u>(Suzuki et al., 2018)</u>, consistent with suppression of neuronal differentiation through enhanced progenitor maintenance at the end of neurogenesis. A plausible scenario is that these maintained progenitors subsequently enter gliogenic pathways, including differentiation into glial-IPC, leading to increased astrocyte output in adults under N2NLB overexpression. To directly examine the fate of dividing cells, we labeled proliferative cells with EdU at P0 and assessed marker expression at P1. A higher proportion of electroporated cells incorporated EdU in the N2NLB-overexpressing population than in controls (Fig. 5D-E), consistent with enhanced maintenance of proliferative progenitors at P1. Among N2NLB-expressing EdU cells, the fraction of Olig2 glial lineage cells was increased relative to controls, whereas NeuN neurons remained at very low levels and were not detectably changed. Together, these data suggest that N2NLB both maintains proliferation and biases dividing progenitors toward glial lineage commitment around birth.

### N2NLB suppresses neuronal genes and activates cell proliferation programs

To elucidate molecular correlates of N2NLB-induced astrogenesis in mice, we performed single-nucleus RNA-seq on FACS-isolated nuclei from N2NLB- or control EGFP-expressing cells collected at P1, P4, and P8 after E15.5 electroporation (Fig. 6A). Integration with a reference dataset of the developing mouse cortex <u>(Di Bella et al., 2021)</u> revealed major neural populations, including neuronal subtypes as well as glial-IPC and astrocytes (Fig. 6B-C). Correlation analysis identified genes whose expression varied with the level (“dosage”) of overexpressed N2NLB. Genes negatively correlated with N2NLB were enriched for neuronal processes such as synapse organization, consistent with the negative correlations observed for endogenous N2NL in the human fetal cortex (Fig. 1C, Table S2). Conversely, genes positively correlated with N2NLB were enriched for metabolic and proliferative pathways, including RNA splicing, intracellular transport, and chromosome condensation (Fig. 6D-E, Table S2). These data indicate that N2NLB expression is associated with suppression of neuronal gene programs and upregulation of proliferation-related and homeostatic programs during gliogenesis.

**Figure 6.**
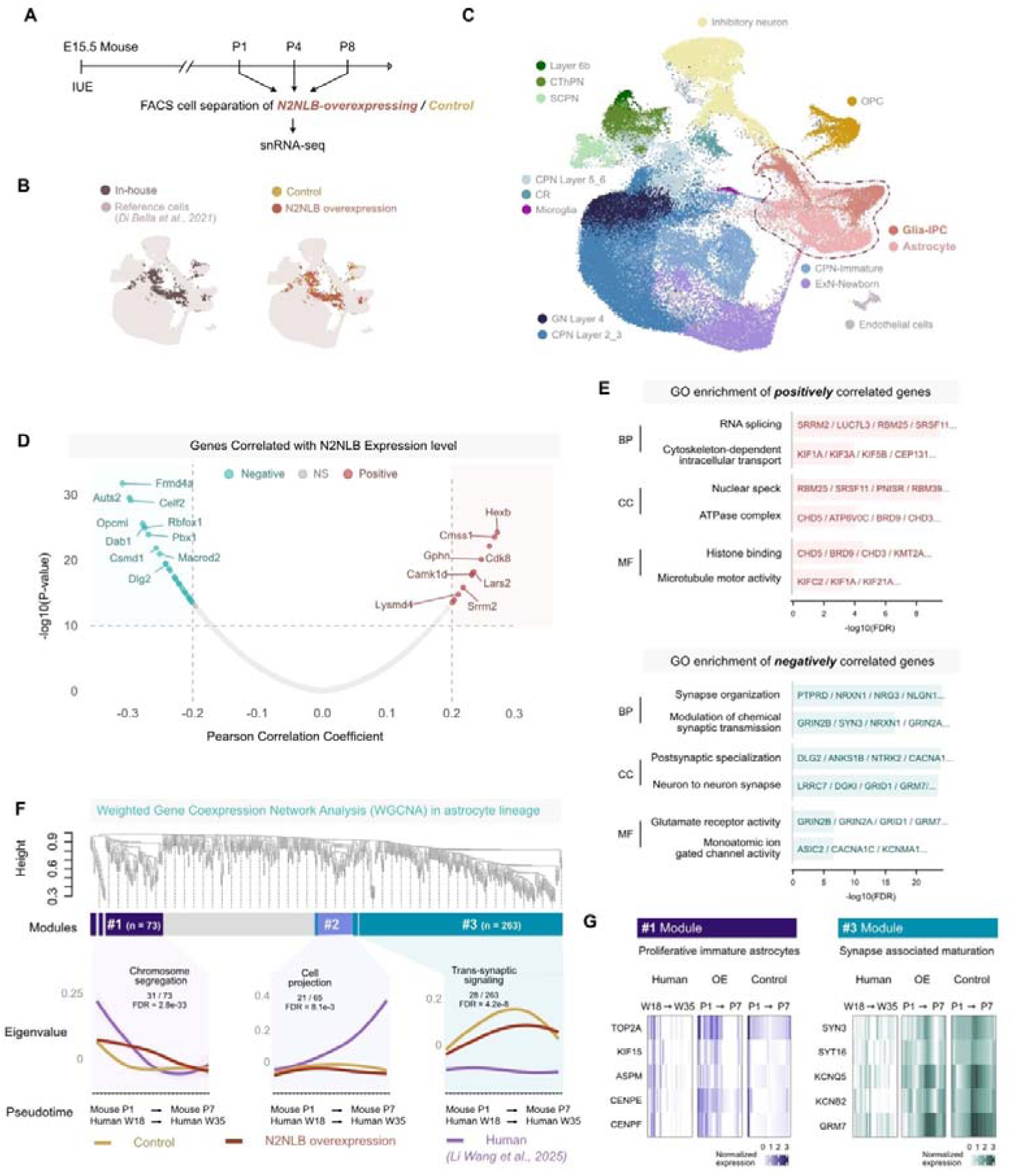
N2NLB overexpression shifts gene expression signature from neuronal state to more proliferative state during mouse gliogenesis. **(A)** Experimental paradigm for single-nucleus RNA-seq following N2NLB overexpression during mouse cortical gliogenesis. Nuclei from N2NLB- or EGFP-expressing cells were isolated at P1, P4, and P8 and subjected to transcriptome profiling. **(B)** Integration of the obtained dataset with a published single-cell transcriptomic reference of the developing mouse cortex <u>(Di Bella et al., 2021)</u>. **(C)** UMAP visualization of the integrated dataset, revealing distinct clusters corresponding to major cortical cell types. **(D)** Genes whose expression levels show positive or negative correlation with the dosage of overexpressed N2NLB. **(E)** Gene Ontology analysis of correlated genes. Notably, genes negatively correlated with N2NLB dosage are enriched for neuronal functions such as synapse organization, whereas genes positively correlated are enriched for proliferative and metabolic processes. **(F)** WGCNA identified gene modules with shared expression dynamics along the astrocyte differentiation trajectory. Module #1 contains genes with higher expression in humans than in mice and associated with cell-cycle regulation, whereas Module #3 contains genes more highly expressed in mice and related to neuronal functions. **(G)** Representative expression dynamics of Module #1 and Module #3 genes during gliogenesis, illustrating their divergent patterns and species differences.

We next asked whether N2NLB shifts mouse transcriptional programs toward patterns enriched in human astrocyte-lineage trajectories. Using cross-species alignment of differentiation trajectories prepared in our previous study <u>(Sheu et al., 2025)</u> together with weighted gene co-expression network analysis (WGCNA)-defined modules along the astrocyte lineage (Fig. 6F, Table S3), we found that Module 1 genes, which show higher expression in humans than in mice and are associated with cell proliferation, were upregulated by N2NLB (Fig. 6G). In contrast, Module 3 genes, which show higher expression in mice and are enriched for neuronal functions (e.g., trans-synaptic signaling), were downregulated by N2NLB. Thus, N2NLB overexpression shifts module-level transcriptional signatures toward a pattern more consistent with human-enriched proliferative programs and away from neuronal programs, providing a mechanistic correlate for enhanced astrocyte-lineage proliferation.

## Discussion

Our single-cell transcriptomic analyses of human fetal cerebral cortices revealed that N2NL is expressed not only in neurogenic RG subtypes but also at substantial levels along differentiation trajectories toward astrocytes. Functional assays in an astrocyte culture system further showed that N2NL is required for robust proliferation and that N2NL overexpression is sufficient to enhance proliferation under these conditions. Consistent with these findings, overexpression of N2NL in mouse neural progenitors *in vivo* significantly increased the proportion of astrocytes among their progeny. This phenotype emerged during the early postnatal period and persisted into adulthood. To elucidate cellular dynamics underlying enhanced astrocyte output, we examined gliogenesis in detail and found evidence that N2NL expands proliferative glial precursor populations around birth, including glial-IPC-like states (Fig. 7). Transcriptomic profiling of N2NL-expressing cells in the mouse brain further revealed suppression of neuron-associated gene programs and concurrent upregulation of genes involved in cell proliferation and cellular homeostasis. Together, these results support a model in which a human lineage-specific emergence of N2NL can tune the balance between neuronal and glial developmental programs (Fig. 7).

**Figure 7.**
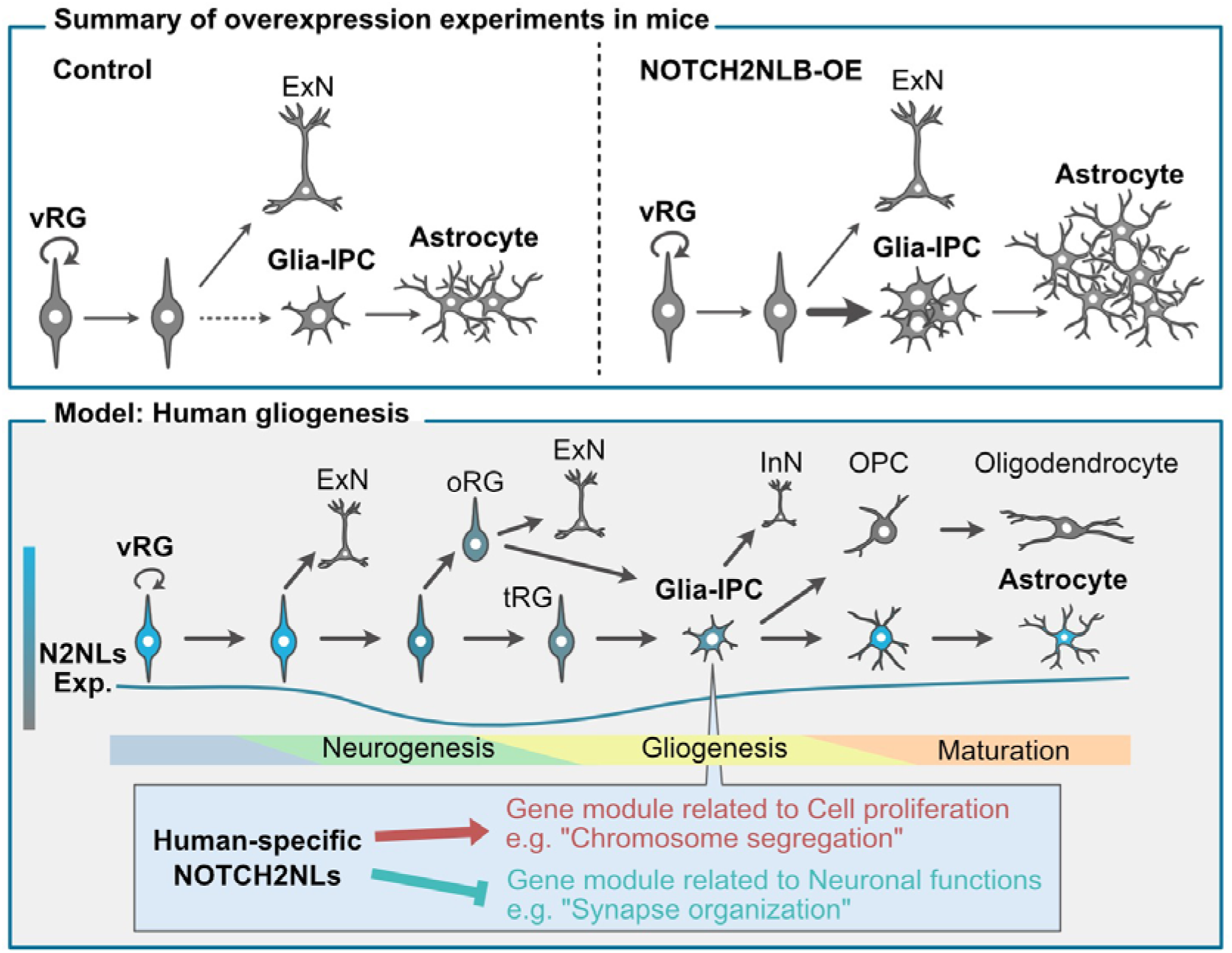
Summary and hypothetical model of human gliogenesis regulated by N2NL. (Top) Summary of the experimental results in this study. NOTCH2NLB enhances glial progenitor program to amplify astocyte output. (Bottom) Hypothetical model of human cortical gliogenesis regulated by human-specific N2NL paralogs. The dynamics of endogenous expression of N2NL paralogs are indicated by the line below the differentiation trajectory and by the color of the cells. InN: inhibitory neurons.

### Mechanistic model: how the human-specific gene N2NL increases astrocyte number

Humans have a substantially higher proportion of astrocytes than other mammals <u>(Allen</u> <u>and Lyons, 2018; Oberheim et al., 2009; Zuchero and Barres, 2015)</u>, yet the mechanisms driving this evolutionary expansion remain largely elusive. Our findings support a model in which N2NL biases RG toward differentiation into glial-IPCs, which subsequently proliferate to expand the astrocyte pool. While lineage progression from fetal RG to mature glia is becoming clearer, current models suggest that mouse RG maintained in an undifferentiated state during neurogenesis eventually gives rise to glial-IPCs and ependymal cells around birth <u>(Li et al., 2021; Spassky et al., 2005; Weng et al., 2019; Zhang et al., 2020)</u>. These glial-IPCs, which express EGFR and Olig2, can differentiate into oligodendrocytes (via OPCs), astrocytes, and inhibitory neurons that selectively migrate to the olfactory bulb <u>(Gao et al., 2025; Li et al., 2021; Zhang et al., 2020)</u>. Alternative routes have also been proposed, such as RG differentiating directly into astrocytes or oligodendrocytes without passing through a glial-IPC stage <u>(Liu et al., 2022; Zhou et al., 2025)</u>. These distinct pathways may correspond to specific RG subtypes: highly proliferative Emx1 /Id3 RG, which preferentially generate gray matter astrocytes via glial-IPCs, and less proliferative Emx1 /Ascl1 RG, which are more likely to produce white matter astrocytes via a glial-IPC-independent route <u>(Zhou et al., 2025)</u>.

In our study, N2NL increased the number of Olig2 glial-IPC-like cells, raising the question of whether it also affects the Emx1 /Ascl1 lineage. Although we observed suppression of neuron-associated genes upon N2NL expression, the precise downstream signaling pathways remain to be clarified. During neurogenesis, N2NL is known to activate Notch signaling to maintain stemness in vRGs <u>(Suzuki et al., 2018)</u>; however, whether the same Notch-dependent mechanism drives the gliogenic effects observed here remains to be directly tested. While Notch signaling is well established as a promoter of astrocyte formation <u>(Gaiano et al., 2000; Namihira et al., 2009; Wang and Barres, 2000)</u>, the specific progenitor stage and cellular sources of Notch ligands during gliogenesis are poorly defined. Recent genetic studies in mice indicate that Notch signaling is essential for astrocyte production specifically during the RG-to-glial-IPC transition, whereas Notch activity at later stages primarily suppresses OPC differentiation without strongly influencing total astrocyte numbers <u>(Gao et al., 2025; Guo et al.,</u> <u>2023)</u>. These observations are consistent with the hypothesis that N2NL may enhance astrocyte production by elevating Notch signaling at the RG-to-glial-IPC transition. Given that Notch signaling represses the proneural gene Ascl1, N2NL may preferentially promote the Ascl1L pathway (Id3L RG → glial-IPC → astrocyte), which is associated with higher proliferative capacity, thereby increasing astrocyte output.

### Comparative considerations: human and primate-specific aspects of astrocyte lineage

Comparative studies highlight both notable differences and key similarities between human and mouse corticogenesis. A defining feature of human cortical development is the diversification of radial glia (RG) into ventricular RG (vRG/tRG) and outer RG (oRG) within the expanded outer subventricular zone <u>(Nowakowski et al., 2016)</u>, a pattern shared with other gyrencephalic mammals such as ferrets and macaques. Evidence from the human fetal cortex suggests that tRG primarily generates gray matter astrocytes, whereas oRG preferentially gives rise to white matter astrocytes <u>(Allen et al., 2022)</u>. Both tRG and oRG can also produce glial-IPCs, which subsequently generate oligodendrocytes, astrocytes, and inhibitory neurons that migrate into the neocortex <u>(Wang et al., 2025)</u>. Notably, human gliogenesis begins while neurogenesis remains active, resulting in substantial temporal overlap between these processes—contrasting with rodents, where the transition from neurogenesis to gliogenesis occurs more abruptly around the perinatal period.

Our analyses indicate that although N2NL expression is relatively modest in late-gestational tRG and oRG, it is upregulated in glial-IPCs and increases further as cells differentiate into astrocytes (Fig. 1). These differences in RG lineage architecture caution against directly extrapolating N2NL functions in human gliogenesis solely from mouse models. Nonetheless, if analogous mechanisms operate in humans, N2NL is well positioned to act during the RG to glial-IPC transition and/or within astrocyte lineage states to increase astrocyte output. Moreover, our finding that N2NL supports proliferation in a human astrocyte culture system suggests that N2NL-dependent signaling, or potentially additional pathways, may contribute not only to early lineage decisions but also to maintenance of astrocyte proliferative competence. Direct functional studies of N2NL in late-gestational human cortex remain limited by tissue accessibility, making human brain organoids an important future avenue for defining the precise cellular contexts and mechanisms by which N2NL regulates human gliogenesis.

### Implications for glioblastoma and glial tumorigenesis

Beyond its role in normal brain development, N2NL appears to be implicated in glioblastoma biology <u>(Funato et al., 2021)</u>. Although glioblastomas do not always originate from mature glial cells, accumulating evidence suggests that glial-IPCs or OPC-like progenitors may serve as cells of origin <u>(Bao et al., 2006; Bhaduri et al., 2020; Chen et al., 2012, 2010; Couturier et al., 2020; Cusulin et al., 2015; Galli et al., 2004; Liu et al., 2011, 2006; Meyer et al.,</u> <u>2015; Singh et al., 2004)</u>. Our finding that N2NL can promote expansion of proliferative glial precursor states suggests that it may contribute to early steps of gliomagenesis in appropriate contexts. Furthermore, glioblastoma tissue exhibits remarkable cellular plasticity, in which differentiated cells can dedifferentiate into a stem-like state, enabling bidirectional conversion between cell types <u>(Neftel et al., 2019)</u>. Within such tumors, high N2NL expression might help maintain or expand stem-like populations. Therefore, understanding how N2NL regulates astrocyte-lineage development may illuminate mechanisms underlying glioblastoma initiation, progression, and therapeutic resistance.

### Human- or primate-specific features of astrocytes

Human astrocytes exhibit unique properties beyond increased number and size <u>(Oberheim Bush and Nedergaard, 2017; Oberheim et al., 2009, 2006)</u>. Compared to murine counterparts, human astrocytes display markedly incomplete tiling, characterized by overlap of processes between neighboring cells. Some astrocytes extend long processes spanning multiple cortical layers and intruding into adjacent territorial domains. Strikingly, these human-specific morphological features are recapitulated when human glial progenitors are transplanted into the mouse brain, indicating that enlarged size and complex process architecture are cell-autonomous properties not overridden by the murine brain environment <u>(Han et al., 2013)</u>. This raises the possibility that human-specific genes contribute to the emergence of these traits. Although N2NL overexpression did not overtly alter astrocyte size in our system, it remains possible that it modulates finer aspects of morphology, such as process overlap, or functional properties such as intercellular coupling and Ca² wave propagation. Future studies will be required to determine whether such human-specific astrocyte features are governed by genetic mechanisms unique to the human lineage, potentially including human-specific genes like N2NL.

## Methods

### Mice

All animal procedures were approved by the Institutional Safety Committee on Recombinant DNA Experiments and the Animal Research Committee of the University of Tokyo. ICR mice were purchased from SLC Japan. Mice were housed in cages with bedding (Avidity Science, TEK-FRESH) and provided with constant access to food (Nippon Crea, Rodent Diet CE-2) and water. The animal facility was maintained at 23 ± 2 °C with 50 ± 10% humidity under a 12-hour light/dark cycle. The day of vaginal plug detection was defined as embryonic day 0.5 (E0.5), and the day of birth was defined as postnatal day 0 (P0). Because sex determination is difficult at embryonic stages, embryos of both sexes were pooled for all analyses.

### Cell lines

Immortalized human astrocytes (Innoprot, Cat# P10251-IM) were cultured on vessels coated with poly-L-lysine hydrobromide (Fujifilm Wako Pure Chemical Corporation, Cat# 166-19081) in Astrocyte Medium (Innoprot, Cat# P60101), according to the manufacturer’s instructions.

### Preparation of lentiviruses

HEK293T cells were seeded at 5.0 × 10 cells per 10-cm dish. After 24 hours, cells were transfected with a plasmid mixture consisting of 3.8 µg pMD2.G, 5.8 µg psPAX2, and 7.62 µg of the transfer vector (pLPGK-H2B-mCherry or pLC-N2NLB-ires-H2B-mCherry). Transfection was performed in 1,250 µL Opti-MEM supplemented with 53.1 µL X-tremeGENE HP DNA Transfection Reagent (Roche). After 24 hours, the medium was replaced, and cells were cultured for an additional 48 hours. Viral supernatants were collected, filtered through a 0.45 µm filter, and concentrated by ultracentrifugation at 50,000 × g for 2 hours at 16 °C using a 20% sucrose/PBS cushion. The viral pellet was resuspended in 100 µL cold PBS, aliquoted (20 µL each), and stored at −80 °C.

### DNA constructs

iOn-N2NLB-ires-EGFP and iOn-N2NL-ires-H2B-mCherry constructs were generated by inserting PCR-amplified N2NLB-ires or H2B-mCherry fragments into iOn vector backbones (Kumamoto et al., 2020) using the In-Fusion HD Cloning Kit (Takara, Cat# CLN639649).

### Lentiviral transduction

To assess the effect of N2NLB overexpression in human astrocyte cultures, lentiviruses encoding pLPGK-H2B-mCherry (control; Ctrl) or pLC-N2NLB-ires-H2B-mCherry (N2NLB overexpression; N2NL) (Suzuki et al., 2018) were transduced. For proliferation analysis, EdU was added to the culture medium 3 days post-infection. After 24 hours of incubation, cells were fixed with 4% paraformaldehyde (PFA).

### Nucleofection of plasmids and siRNAs

Human astrocyte cultures were dissociated and resuspended in 100 µL Nucleofector Solution from the P3 Primary Cell 4D-Nucleofector™ X Kit L (Lonza, Cat# V4XP-3024). A total of 1 × 10 cells were mixed with 1 µg pLPGK-H2B-mCherry plasmid. For gene knockdown experiments, 200 nM siRNA targeting N2NL (Qiagen, FlexiTube GeneSolution GS388677 [#1027416]) or AllStars Negative Control siRNA (#1027281) was co-transfected. Electroporation was performed using the DS-150 program on the 4D-Nucleofector™ X Unit (Lonza). Transfected cells were seeded onto poly-L-lysine–coated 8-well chamber slides. For proliferation assays, EdU was added to the culture medium 3 days post-transfection, followed by fixation with 4% PFA for 15 minutes at room temperature after 24 hours of incubation.

### RNAscope in situ hybridization

Human astrocytes were cultured on poly-L-lysine coated 8-well chamber slides and fixed with 4% PFA. RNA in situ hybridization was performed using the RNAscope Multiplex Fluorescent Detection Kit v2 (Advanced Cell Diagnostics, Cat# 32310) according to the manufacturer’s instructions (User Manual UM323110). A probe specific for N2NLB (ACD, Cat# 1076651-C2) was used. Images were analyzed using ImageJ (NIH), and the number of fluorescent puncta per cell was quantified to assess gene expression levels.

### In utero electroporation

*In utero* electroporation was performed with modifications to established protocols (Tabata and Nakajima, 2001; Yamauchi et al., 2025). Timed-pregnant mice at E15.5 were anesthetized with isoflurane. Plasmid solutions (1-2 mg/mL DNA) were injected into the embryonic lateral ventricles using heat-pulled glass capillaries. For control experiments, a plasmid mixture containing CAG-hyPBase (1.0 µg/µL), iOn-CAG-ires-mCherry (0.2 µg/µL), and iOn-CAG-ires-EGFP (0.2 µg/µL) was injected. For N2NLB overexpression, iOn-CAG-N2NLB-ires-EGFP (0.2 µg/µL) was used in place of iOn-CAG-ires-EGFP. Electroporation was performed using tweezer electrodes (Nepa Gene, Cat# CUY650P5) connected to a NEPA21 Type II electroporator (Nepa Gene) with the following parameters: 35-50 V, 50 ms pulse duration, 950 ms interval, and 4-5 pulses. Embryos were returned to the abdominal cavity, and dams were sutured and allowed to recover on a heating plate. Brains were collected at E18.5, P1, P4, P8, P15, and P56.

### Histology and immunostaining

Immunostaining was performed as described previously(Suzuki et al., 2018, 2012). Mice were perfused transcardially with ice-cold PBS followed by 4% PFA in PBS. Brains were dissected, post-fixed overnight at 4 °C, and sectioned into 100 µm slices using a vibratome (Leica, VT1000S). Sections were washed in PBS containing 0.1% Triton X-100 (PBST) and blocked in PBS containing 0.3% Triton X-100 and 3% donkey serum for 30 minutes. Sections were incubated with primary antibodies (Table S4) for 1–2 days at 4 °C, followed by incubation with secondary antibodies (Table S4) for 2-4 hours at room temperature. Sections were mounted using DAKO Glycerol Mounting Medium (Agilent, Cat# C0563). Imaging was performed using an FV3000 confocal microscope (Olympus), and images were processed with Fiji/ImageJ (Schindelin et al., 2015).

### Quantification of neuron-glia ratio

Brains collected at P4, P8, P15, and P56 were processed for immunohistochemistry as described above. Cell identities were defined based on morphology and cortical localization. Neurons were defined as pyramidal excitatory neurons located in layers 2/3, characterized by large somata and prominent apical dendrites. Glial cells were defined as cells distributed throughout the cortex with small somata and more than five radial processes. Cells not fitting these criteria, mainly located near the ventricles, were classified as “others.” Quantification was performed on the same sections for mCherry-only-expressing cells (internal control) and EGFP-expressing cells (experimental group). The glia ratio was calculated as Glia / (Neurons + Glia).

### Morphological analysis of astrocytes

Mice electroporated at E15.5 were perfused at P56. High-magnification Z-stack images of individual cortical astrocytes were acquired using an FV3000 confocal microscope (Olympus) with a 20× objective lens. mCherry-positive/EGFP-negative cells were analyzed as controls, whereas EGFP-positive/mCherry-negative cells were analyzed as the N2NLB overexpression group. Astrocytic identity was confirmed by Sox9 immunostaining. Z-stack images were projected using maximum-intensity projection in ImageJ (NIH), and cell area was measured by manually outlining the fluorescence-defined cell morphology.

### EdU proliferation assay

Cell proliferation was assessed using 5-ethynyl-2′-deoxyuridine (EdU; Fujifilm, Cat# 052-08843), administered 24 hours before fixation. For in vivo experiments, EdU (5 µg/g body weight) was injected intraperitoneally into pregnant dams at E17 or subcutaneously into pups at P0. For in vitro experiments, EdU was added to culture media at a final concentration of 10 µM. EdU detection was performed using the Click-iT EdU Alexa Fluor 647 Imaging Kit (Thermo Fisher Scientific, Cat# C10340) according to the manufacturer’s instructions.

### Single cell transcriptomics

#### Sample preparation and sequencing

To analyze transcriptomic profiles of transfected cells, in utero electroporation was performed at E15.5 using a plasmid mixture containing CAG-hyPBase (1.0 µg/µL), iOn-CAG-ires-mCherry (0.2 µg/µL), and iOn-CAG-N2NLB-ires-EGFP (0.2 µg/µL). Brains were collected at P1, P4, and P8 and immediately frozen in liquid nitrogen. Nuclei were isolated using a combination of mechanical homogenization and density gradient centrifugation as previously described (Frey et al., 2023). Frozen tissues were thawed and homogenized in 54% Percoll buffer using sequential passage through 23G and 27G needles. The homogenate was incubated with 10% NP-40 on ice for 15 minutes. Nuclei were purified by centrifugation at 20,000 × g for 10 minutes at 4 °C through a Percoll density gradient composed of 31% and 35% Percoll layers.

Fluorescence-activated nuclei sorting was performed using a BD FACS Melody sorter (BD Biosciences). GFP-positive nuclei were collected as the N2NL overexpression group, whereas nuclei expressing mCherry but not GFP (mCherryD/GFPD) were collected as the internal control group. Sorted nuclei were collected into tubes containing STEM-CELLBANKER GMP grade (Takara, Cat# 11924). Library preparation and sequencing were performed with the assistance of 2024PAGS. Libraries were prepared using the 10x Chromium GEM-X Single Cell 3′ Kit v4, and sequencing was carried out on an Element AVITI platform.

### Quantification of overexpressed N2NLB transcript in single-nuclei RNA sequencing

To enable detection of transcripts derived from the N2NLB overexpression construct, the insert sequence of the N2NLB plasmid was added to both the reference FASTA file and the corresponding GTF annotation file. The mouse reference genome (GRCm39 FASTA) and GTF annotation file (version 2024-A) were obtained from the Cell Ranger Download Center. Sequencing reads were processed using Cell Ranger v9.0.1 with the modified FASTA and GTF files to generate gene-barcode count matrices, allowing accurate quantification of transgenic N2NLB expression.

### Integration and clustering

Downstream single-cell RNA sequencing analyses were performed using Seurat v5 (Satija et al., 2015; Stuart et al., 2019). The in-house postnatal mouse N2NL overexpression/control dataset was integrated with a previously published mouse cortical development dataset (Di Bella et al., 2021), selecting developmental stages from E16 to P4 as an external reference.

Quality control filtering was applied to retain high-quality cells based on the following criteria: unique molecular identifier counts (nCount_RNA) > 1,000, gene counts (nFeature_RNA) > 1,000, log10(genes per UMI) > 0.8, and mitochondrial gene percentage (percent.mt) < 7.5%. Data were initially split into separate Seurat objects corresponding to individual batches or mouse samples. Each object was normalized and scaled using SCTransform (Hafemeister and Satija, 2019), and technical covariates (nCount_RNA, nFeature_RNA, percent.mt, and CC.Difference) were regressed out to minimize effects from sequencing depth, transcript complexity, mitochondrial content, and cell cycle state.

Highly variable genes (HVGs) were identified independently for each batch. To reduce bias during integration across heterogeneous developmental stages, samples were grouped into three transcriptomically similar temporal windows (early: E16-E17; middle: E18-P1; late: P3-P8). HVGs were selected within each window using SelectIntegrationFeatures, and the union of these genes was used for downstream integration. Batch correction was performed in reduced dimensional space using reciprocal principal component analysis (RPCA) (Stuart et al., 2019). Principal component analysis was conducted on the scaled data, and the first 30 principal components were retained based on elbow plot inspection. Uniform manifold approximation and projection (UMAP) visualization was generated using default parameters applied to the integrated embeddings. Clustering was performed using the Louvain algorithm with k-nearest neighbors set to 25 and a resolution parameter of 1.2. Cluster identities were assigned based on the top 50 differentially expressed genes per cluster, identified using a Wilcoxon rank-sum test with Bonferroni correction, ranked by −log10(adjusted value), and filtered to require >30% expression within the cluster and a log2 fold change > 0.5, together with known canonical marker genes.

### Pseudotime analysis

To reduce potential biases arising from unequal numbers of cells across developmental stages, cells from each stage were aggregated into an equal number of “metacells” (n = 30 per stage). Metacells were generated by constructing a shared nearest neighbor (SNN) graph based on k-nearest neighbors using the Louvain algorithm (Waltman and van Eck, 2013), and gene expression values for each metacell were calculated as the average expression across constituent cells.

Pseudotime inference was performed using L1-penalized ordinal regression as implemented in the psupertime R package (Macnair et al., 2022). Ordinal labels corresponding to developmental stages were supplied to the model, which projects gene expression profiles onto a continuous pseudotime axis. Model training was conducted using 10-fold cross-validation to select the optimal regularization parameter, and the best-performing model was used to predict pseudotime values for individual cells.

### Human neurodevelopmental reference and DDRTree trajectory analysis

To provide a human neurodevelopmental reference, publicly available single-cell transcriptomic datasets of human cortical development were integrated into the analysis (Wang et al., 2025). Developmental stages corresponding to estimated post-conception days 54 to 246 (gestational weeks 10-37) were selected to focus on the neurodevelopmental period. Gene expression values were obtained from the SCTransform-normalized assay, in which effects from sequencing depth, batch variation, mitochondrial and ribosomal gene content, and cell cycle differences had been regressed out.

Trajectory analysis was performed using Monocle 2 (Qiu et al., 2017) to construct a discriminative dimensionality reduction tree (DDRTree). Ordering genes were selected based on differential expression analysis between annotated cell types, selecting the top 1,500 genes with values < 0.05. Dimensionality reduction was performed using DDRTree with a maximum of two components, applying reverse graph embedding to learn a principal graph. Parameters were set as follows: sigma = 1e-4, lambda = 10 × number of cells, ncenter = 200, param.gamma = 120, and maxIter = 200. The resulting minimum spanning tree was used to order cells along the trajectory, with pseudotime and state assignments refined by rooting the trajectory at the ventricular radial glia (vRG)-enriched state and defining branches using shortest-path distances and depth-first search. Normalized gene expression patterns were visualized along the inferred DDRTree trajectory.

### Weighted gene co-expression network analysis (WGCNA)

Weighted gene co-expression network analysis (WGCNA) (Langfelder and Horvath, 2008) was performed on a pseudotime-aligned expression matrix to identify gene modules with distinct temporal expression dynamics. To enable direct comparison between mouse and human datasets, gene expression matrices were constructed using a standardized pseudotime grid divided into 30 bins scaled from 0 to 1, rather than using absolute developmental stages. Prior to network construction, gene expression values were z-score normalized to a mean of zero and a standard deviation of one.

The soft-thresholding power was determined using the *pickSoftThreshold* function by evaluating candidate powers from 1 to 20 and selecting the value that best approximated a scale-free topology. Co-expression networks and gene modules were constructed using the *blockwiseModules* function with a signed “Nowick 2” topological overlap matrix (TOM) to allow detection of anti-correlated expression patterns. The minimum module size was set to 30 genes, and modules with similar expression profiles were merged using a merge cut height of 0.1. Module eigengenes were calculated to summarize the expression dynamics of each identified module.

## Data and resource availability

All animals and unique/stable reagents generated in this study are available from the lead contact with a completed Materials Transfer Agreement. All data and the codes for analyzing the data in this study are available from the lead contact. Single cell RNA sequencing data of electroporated cortical cells are available in DDBJ Sequence Read Archive (DRA021462).

## Supplementary tables and figures

### Table supplements

Supplementary table 1: N2NL correlated genes in human cortical development

Supplementary table 2: N2NL correlated genes in mouse in-house datasets

Supplementary table 3: WGCNA module genes and GO term

Supplementary table 4: Antibody list

### Figure supplements

**Figure supplement 1.**
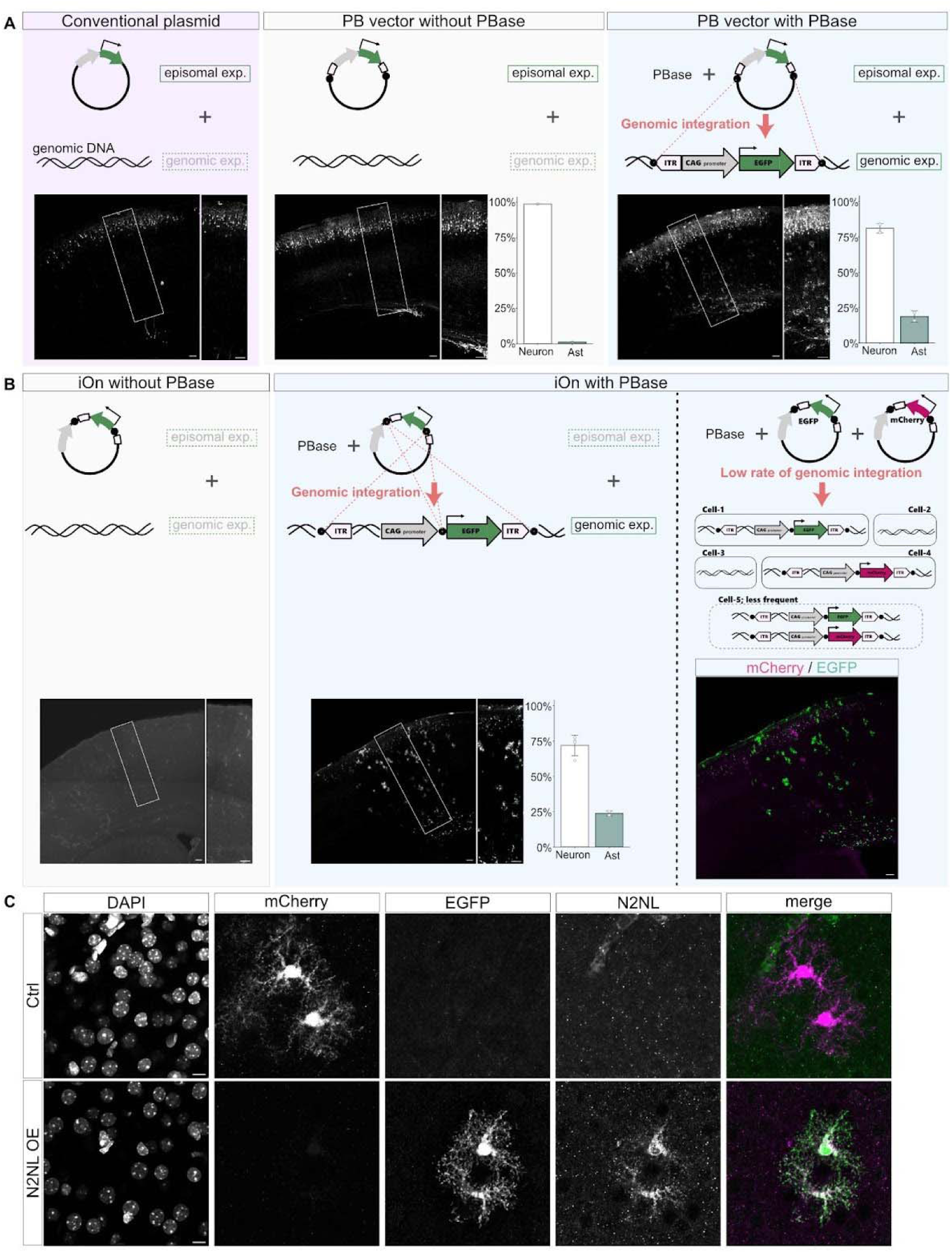
Validation of gene delivery systems. **(A-B)** Comparison of gene-delivery strategies in the developing mouse cortex. (**A**) Episomal plasmid expression is detected primarily in postmitotic neurons generated shortly after IUE, because episomal transgenes are rapidly diluted during progenitor cell divisions. In contrast, genomic integration via the piggyBac transposase (PBase) enables stable and persistent transgene expression in proliferative lineages, including glial cells identifiable by their characteristic morphology. (**B**) The iOn-switch system, which restricts transgene expression to PBase-mediated genomic integration events, yields no detectable episomal expression. Because genomic integration occurs at a low and stochastic frequency, transgenes are sparsely introduced into host cells. As a result, even when multiple constructs are co-electroporated, most labeled cells express only a single transgene, and only a very small fraction co-express multiple constructs. **(C)** N2NLB protein is detected in EGFP-positive cells in the cortex following iOn-switch–mediated overexpression at E15.5. Scale bars: 100 μm (**A**, **B**) and 10 μm (**C**).

**Figure supplement 2.**
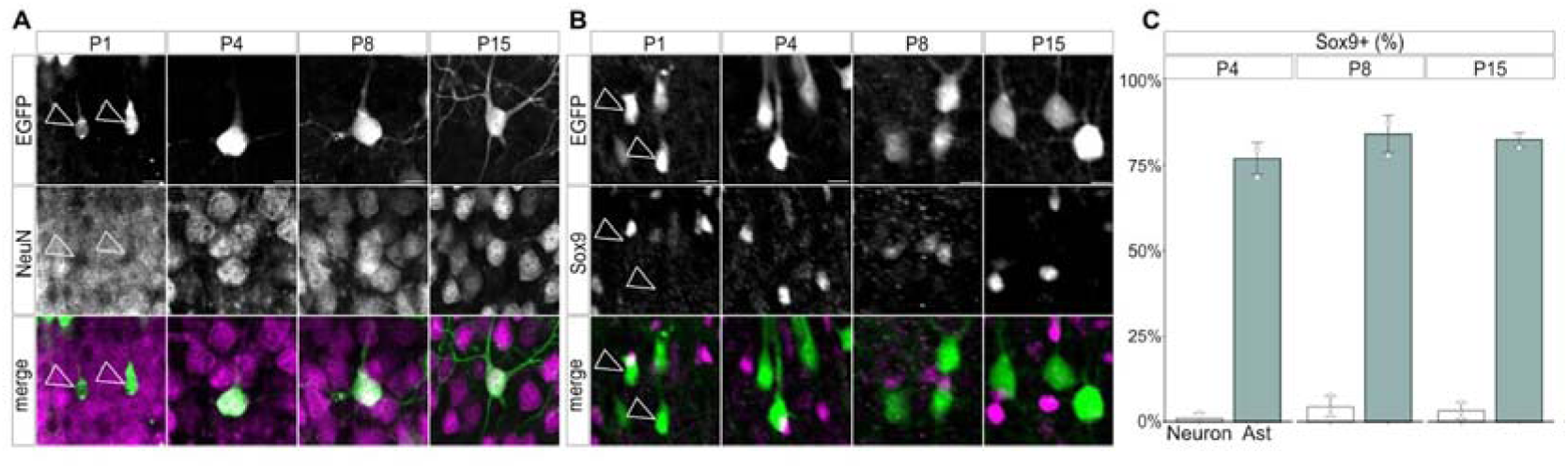
Validation of morphological cell classification using specific molecular markers. **(A–B)** Representative images of electroporated cerebral cortices immunostained for the neuronal marker NeuN (**A**) and the astrocytic marker Sox9 (**B**). At P4, P8, and P15, cells classified as neurons or astrocytes based on morphological criteria are consistently positive for NeuN (**A**, open arrowheads) and Sox9 (**B**, open arrowheads), respectively. In contrast, at P1, cells exhibiting neuron-like morphology are negative for these mature lineage markers, indicating that morphology-based cell-type classification is less reliable at this early postnatal stage due to cellular immaturity. **(C)** Quantification of the proportion of Sox9-positive cells among morphologically defined neurons and astrocytes at P4, P8, and P15. The analysis demonstrates that the vast majority of morphologically defined astrocytes express Sox9, whereas Sox9 labeling is rarely observed in morphologically defined neurons (n = 3 slices per stage). Scale bars, 10 µm.

**Figure supplement 3.**
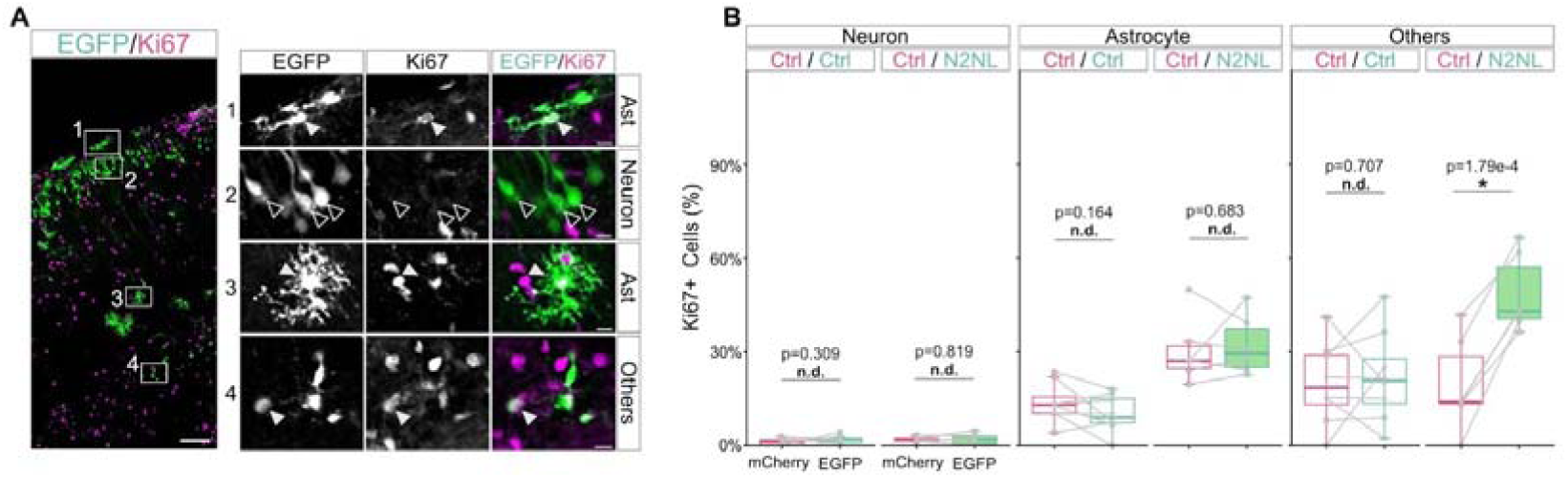
Cell type-specific proliferation index in the P4 cortex. Representative confocal images (**A**) and quantitative analysis (**B**) of Ki67-positive proliferating cells among morphologically defined cell types in the P4 cortex. Mouse cortices were electroporated at E15.5 and analyzed at P4. (**A**) GFP-positive electroporated cells were classified into astrocytes (Ast), neurons, and “others” based on morphological criteria. Filled arrowheads indicate Ki67-positive proliferating cells (double-positive for GFP and Ki67), whereas open arrowheads indicate Ki67-negative, non-proliferating cells. (**B**) Quantification of the percentage of Ki67-positive cells within each morphologically defined cell type. The “others” category comprises cells that do not exhibit clear glial or neuronal morphology and is interpreted as a population of immature glial precursor cells. N2NLB overexpression significantly increased the proportion of Ki67-positive cells specifically within the “others” population (Ctrl/Ctrl: n = 8 slices; Ctrl/N2NL: n = 6 slices). Statistical analysis was performed using a paired t-test. p < 0.05; n.d., not significant. Scale bars, 100 µm (low magnification) and 10 µm (high magnification).

**Figure supplement 4.**
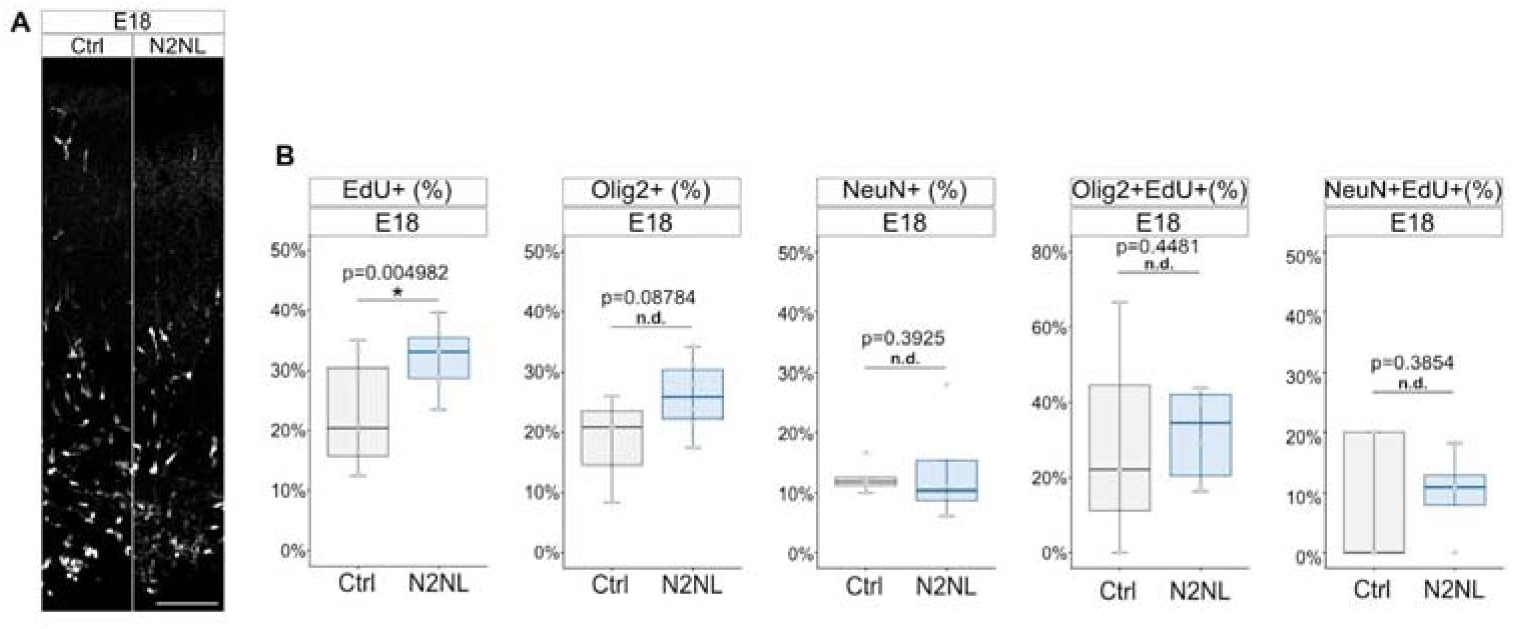
N2NLB promotes cell proliferation but does not significantly change the marker expression at E18.5. **(A)** Representative images of mouse cortices electroporated at E15.5 and analyzed at E18.5. **(B)** Quantification of cell proliferation and fates. Percentages of EdU^+^, Olig2^+^, and NeuN^+^ cells among the total GFP^+^ population. N2NLB significantly increases the proportion of EdU^+^ cells (Ctrl: n = 8 slices; N2NL: n = 10 slices), while Olig2^+^ (Ctrl: n = 3 slices; N2NL: n = 6 slices) and NeuN^+^ (Ctrl: n = 5 slices; N2NL: n = 4 slices) frequencies are not significantly altered. Percentages of Olig2^+^ and NeuN^+^ cells within the EdU^+^ proliferating population (GFP^+^EdU^+^). No significant changes were observed in lineage allocation among dividing cells (Olig2^+^EdU^+^: Ctrl: n = 3 slices; N2NL: n = 6 slices; NeuN^+^EdU^+^: Ctrl: n = 5 slices; N2NL: n = 4 slices). Statistical analyses: Unpaired Student’s t-test (*p < 0.05; n.d., no significant difference). Scale bar: 100 μm.

**Figure supplement 5.**
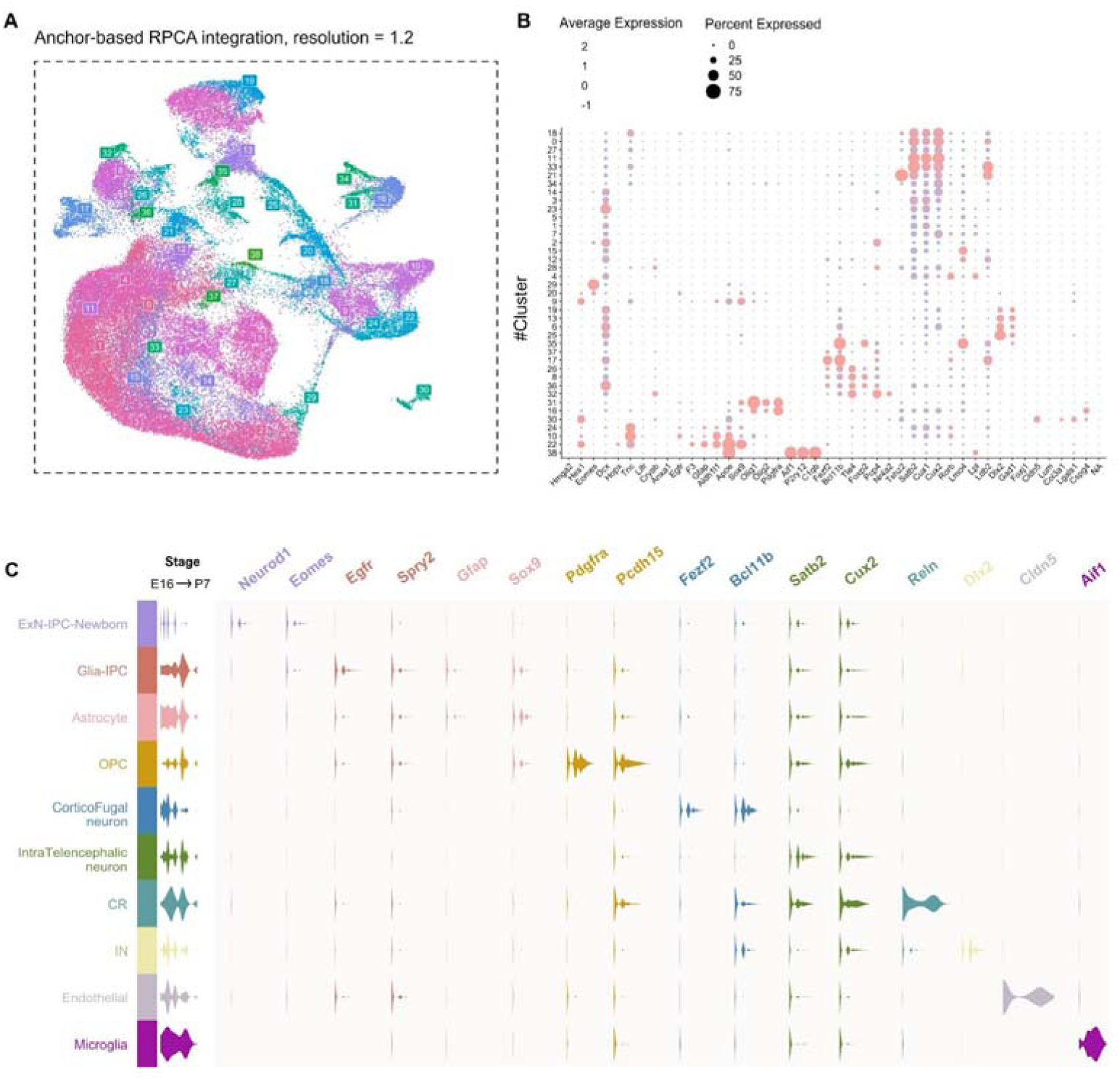
Validation of marker genes in the scRNAseq dataset of human fetal corticogenesis. **(A)** UMAP visualization of the integrated scRNA-seq dataset generated using anchor-based reciprocal PCA (RPCA) integration (clustering resolution = 1.2). Cells are colored according to cluster identity. **(B)** Dot plot displaying the expression patterns of representative marker genes across the identified clusters. Dot size represents the proportion of cells within each cluster expressing the indicated gene, while color intensity denotes the average expression level. **(C)** Stacked violin plots illustrating the expression distributions of cell type–specific marker genes used for cluster annotation. Clusters were assigned to major cell types based on established marker gene expression, including Neurod1 and Eomes for excitatory neuron–intermediate progenitor/newborn neuron populations (ExN-IPC-Newborn); Egfr, Spry2, Gfap, and Sox9 for glial intermediate progenitors and astrocytes (Glia-IPC and Astrocyte); Pdgfra and Pcdh15 for oligodendrocyte precursor cells (OPCs); Fezf2, Bcl11b, Satb2, and Cux2 for excitatory neurons; Reln for Cajal–Retzius cells; Dlx2 for interneurons; Cldn5 for endothelial cells; and Aif1 for microglia.

**Figure supplement 6.**
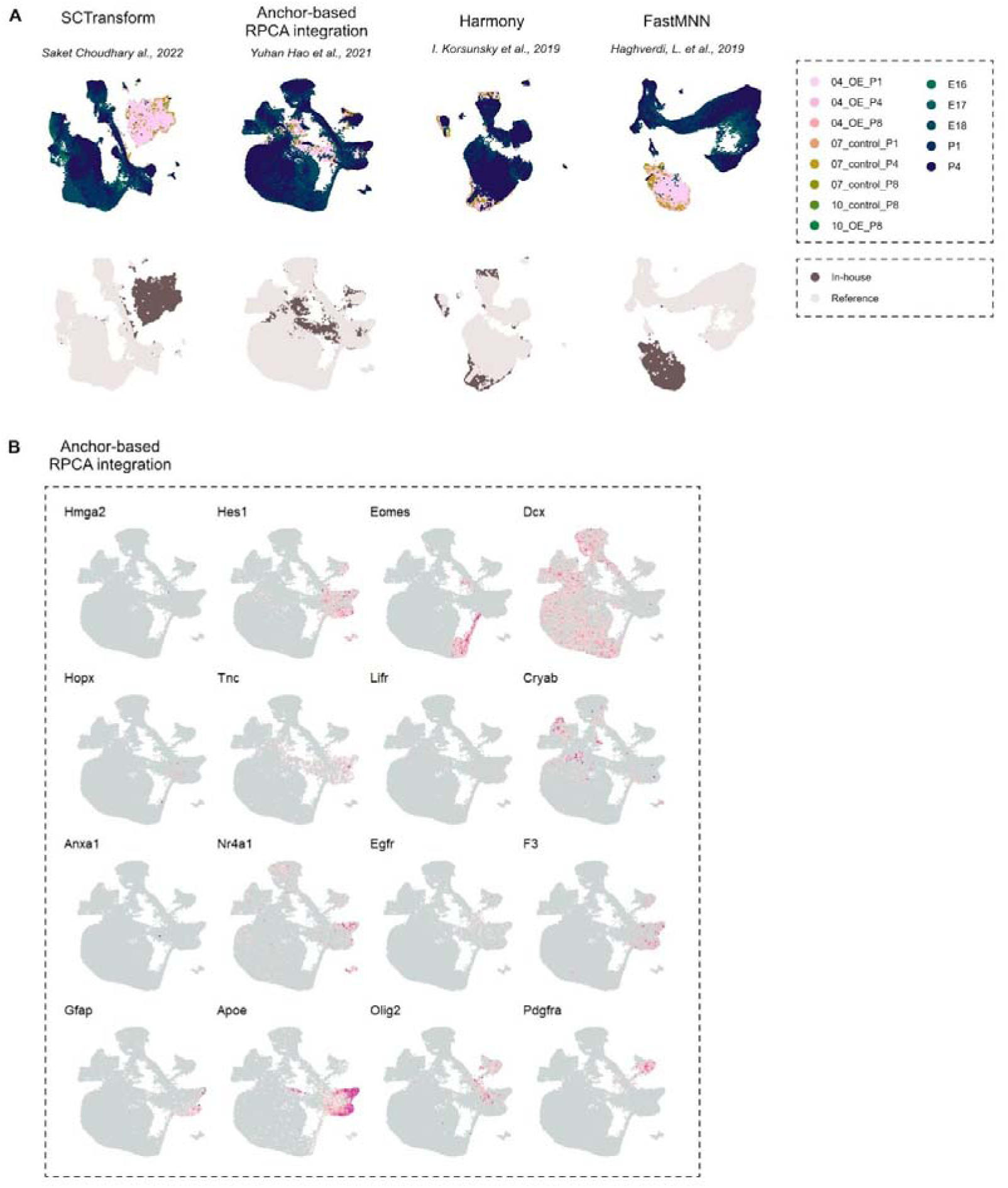
Optimization of data integration and validation of cell type markers for snRNA-seq dataset obtained in house and reference scRNAseq data of mouse corticogenesis. **(A)** Comparison of four single-cell RNA-seq data integration methods—SCTransform, anchor-based reciprocal PCA (RPCA), Harmony, and FastMNN—for integrating the in-house dataset (P1, P4, and P8) with an external developmental reference dataset (Di Bella et al., 2021; E16 to P4). Top panels show UMAP visualizations colored by sample identity and developmental stage. Bottom panels show UMAPs colored by dataset origin, illustrating the degree of mixing between in-house (brown) and reference (beige) cells. Anchor-based RPCA integration was selected for all downstream analyses because it provided the best balance between effective batch mixing and preservation of biologically meaningful structure. **(B)** Feature plots displaying the expression of canonical marker genes projected onto the RPCA-integrated UMAP coordinates. Color intensity indicates relative expression levels. These plots validate the identities of major cell populations, including progenitor cells (Hmga2, Hes1), intermediate progenitors and immature neurons (Eomes, Dcx), astrocyte-lineage cells (Hopx, Tnc, Lifr, Cryab, Anxa1, Nr4a1, Egfr, F3, Gfap, Apoe), and oligodendrocyte-lineage cells (Olig2, Pdgfra).

**Figure supplement 7.**
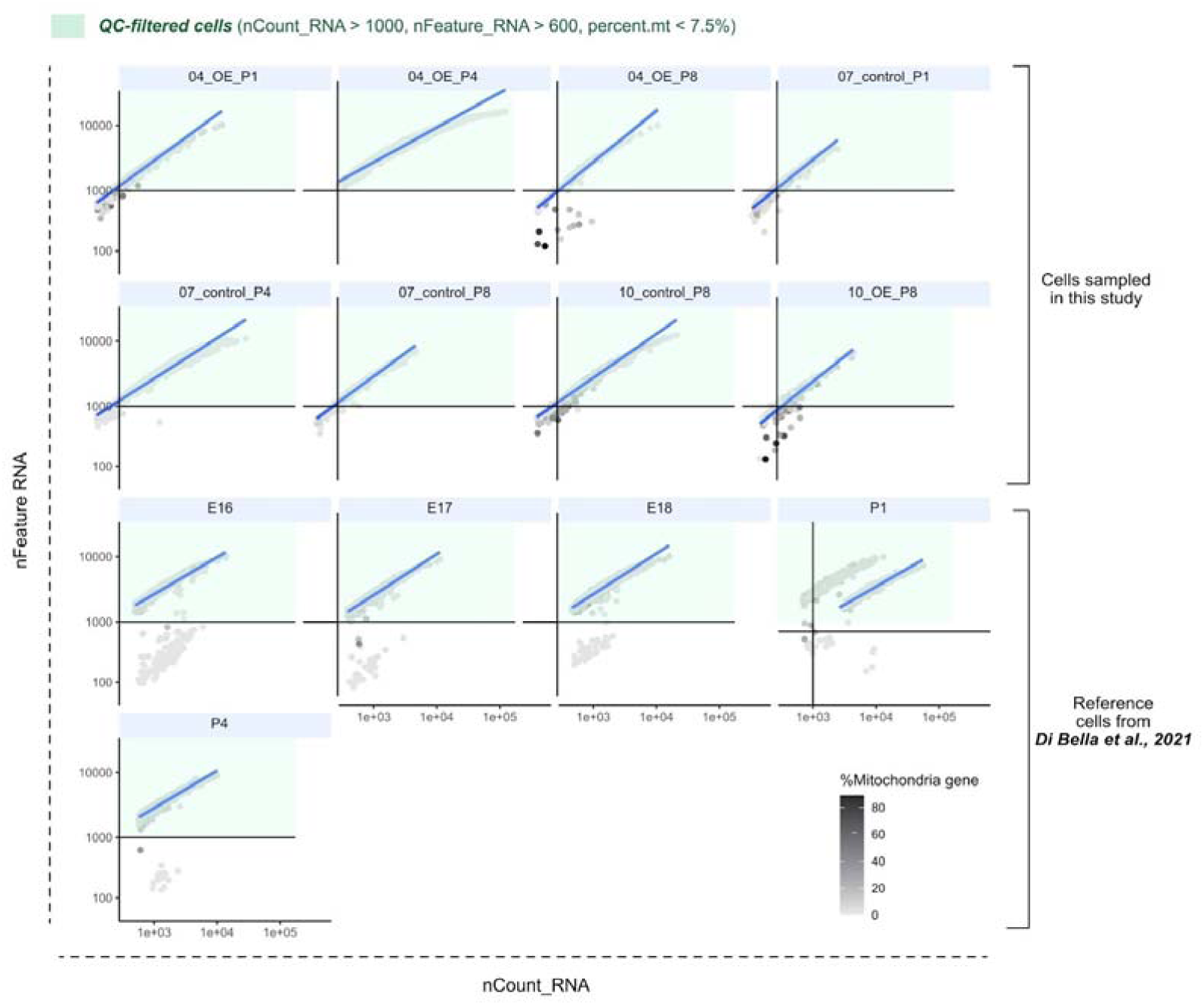
Quality control of snRNA-seq data. Scatter plots showing the relationship between sequencing depth, measured as unique molecular identifier (UMI) counts (nCount_RNA), and the number of detected genes (nFeature_RNA) for each sample. **Top panels** show in-house single-nucleus RNA-seq samples generated in this study at P1, P4, and P8. **Bottom panels** show reference samples from the developing mouse cortex (Di Bella et al., 2021). Cells are colored according to the percentage of mitochondrial gene expression. Green shaded regions indicate high-quality nuclei retained for downstream analyses based on the applied quality-control thresholds: nCount_RNA > 1,000, nFeature_RNA > 600, and mitochondrial gene content < 7.5%.

## Acknowledgements

We are grateful to our colleagues in the lab and common laboratory facilities in the WINGS-LST graduate program at the University of Tokyo. We thank Drs. Tadashi Nomura, Carina Hanashima, Akira Uematsu, Kenji Shimamura and Chiaki Ohtaka-Maruyama for insightful suggestions for the project. Computations were partially performed on the NIG supercomputer at ROIS National Institute of Genetics. This work was supported by JSPS KAKENHI Grant Number JP22H04925 (PAGS). This work was funded by AMED JP25gm7010016h0001, JP24tm0524007 and JP20gm6310006, JST FOREST JPMJFR214T, KAKENHI JP22H02628, JP20H04860 and JP25K02280, MBSJ Tomizawa Jun-ichi & Keiko Fund of Molecular Biology Society of Japan for Young Scientist, Takeda Science Foundation, and SECOM Science and Technology Foundation. R.I, is supported by SPRING GX and X. D. S., is supported by WINGS-LST.

## Additional information

### Author contributions

I. K. S. co-ordinated the project and helped to design, perform and interpret experiments. R. I and I. K. S. designed and performed the experiments and interpreted the results with the help of X. D. S., P. R. and R. A., Y. Y. Y., T. K., and Y. K.. X. D. S. and R. A. performed the analysis of scRNAseq data. K. E. contributed to supervision. R. I., X. D. S., and I. K. S. analyzed the data and prepared figures. R. I., X. D. S., and I. K. S. wrote the paper. All authors read and approved the manuscript.

