## Supplementary figures and images for "Human-specific NOTCH2NL promotes astrogenesis by expanding proliferative glial progenitor states"

### FigS1

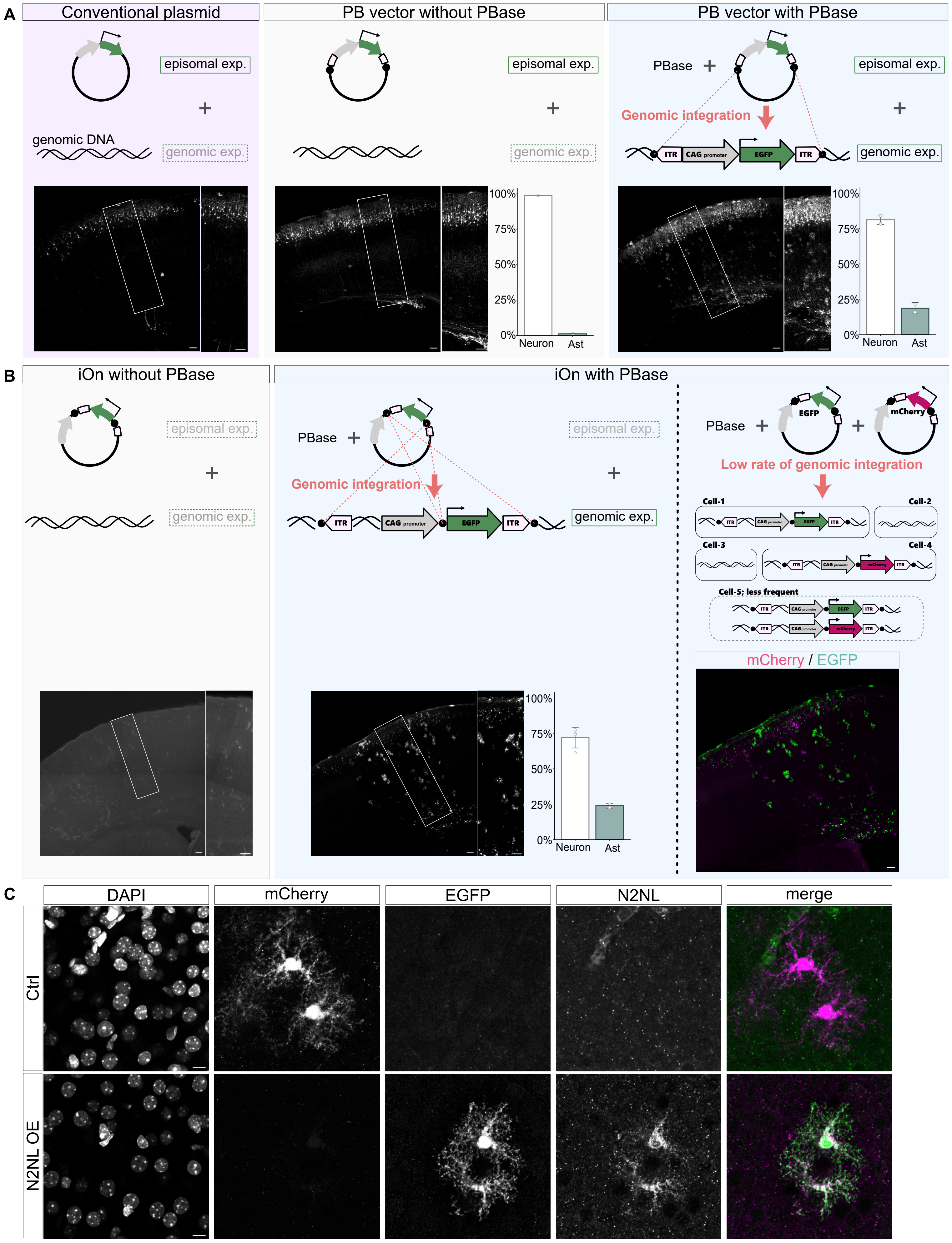

### FigS2

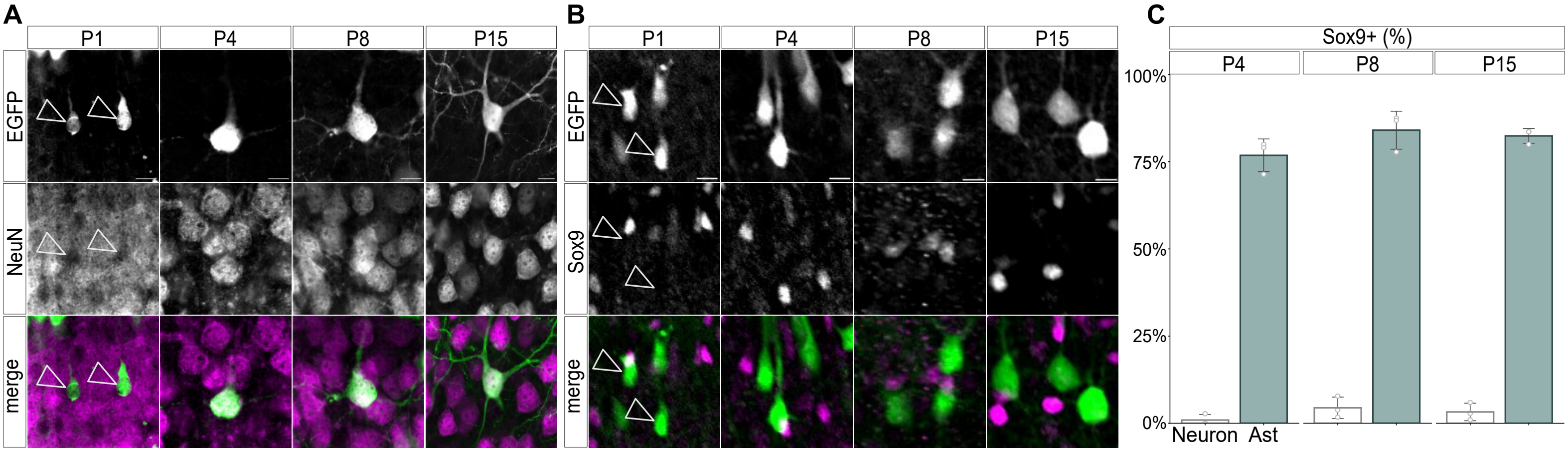

### FigS3

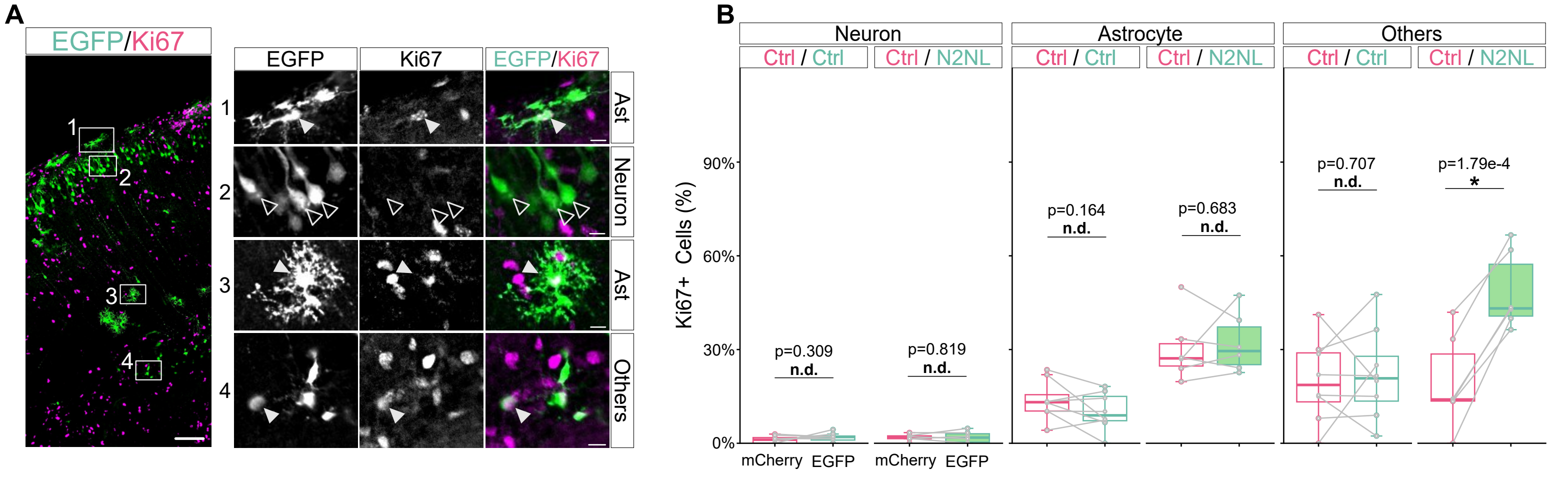

### FigS4

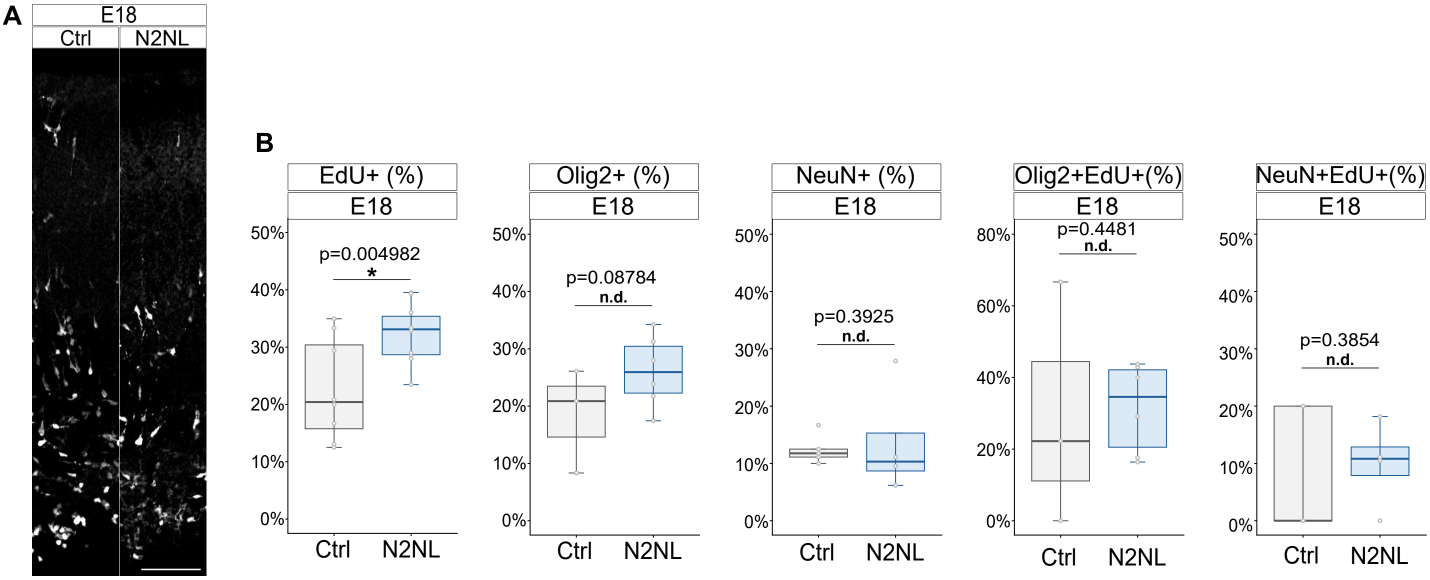

### FigS5

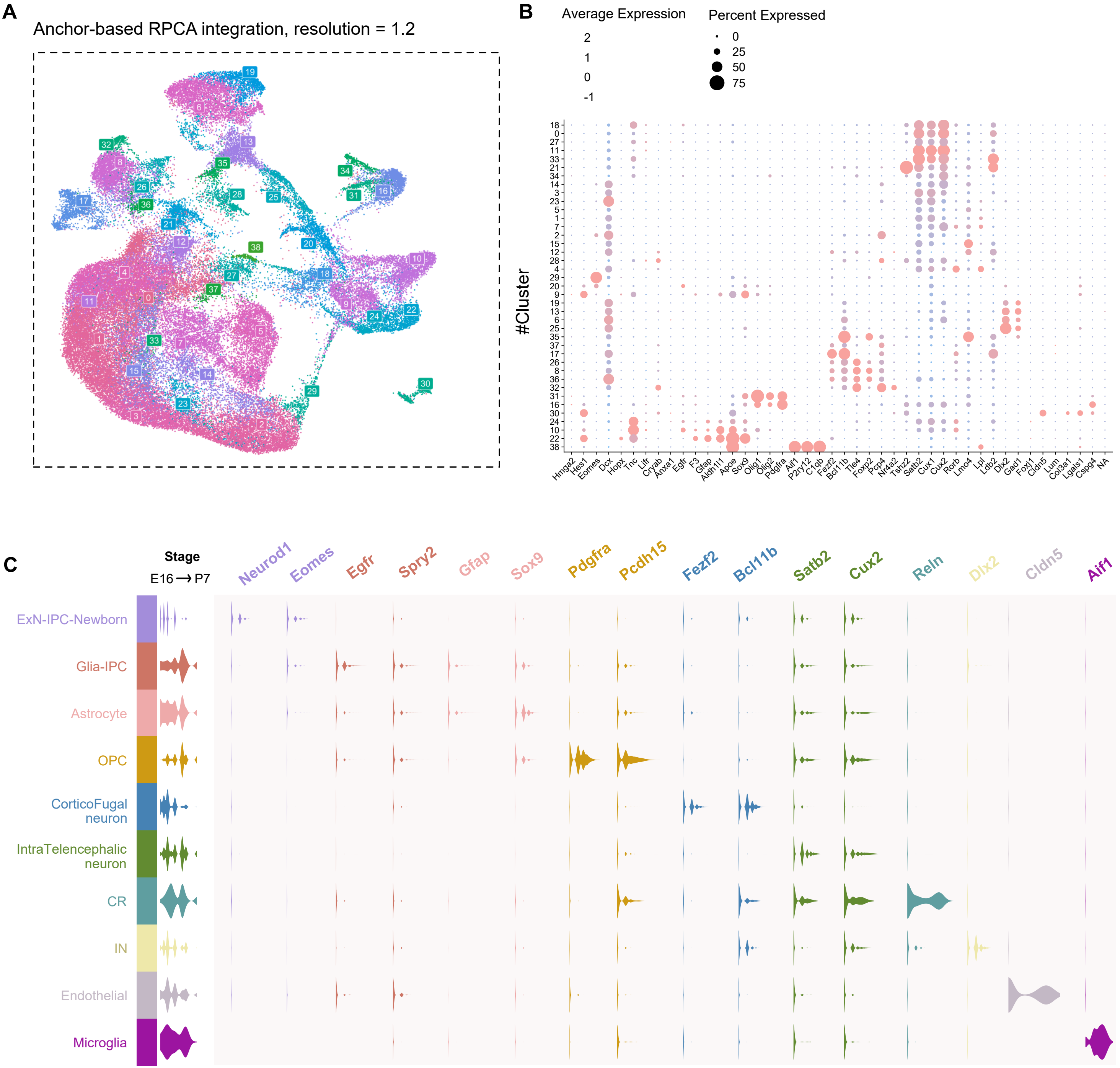

### FigS6

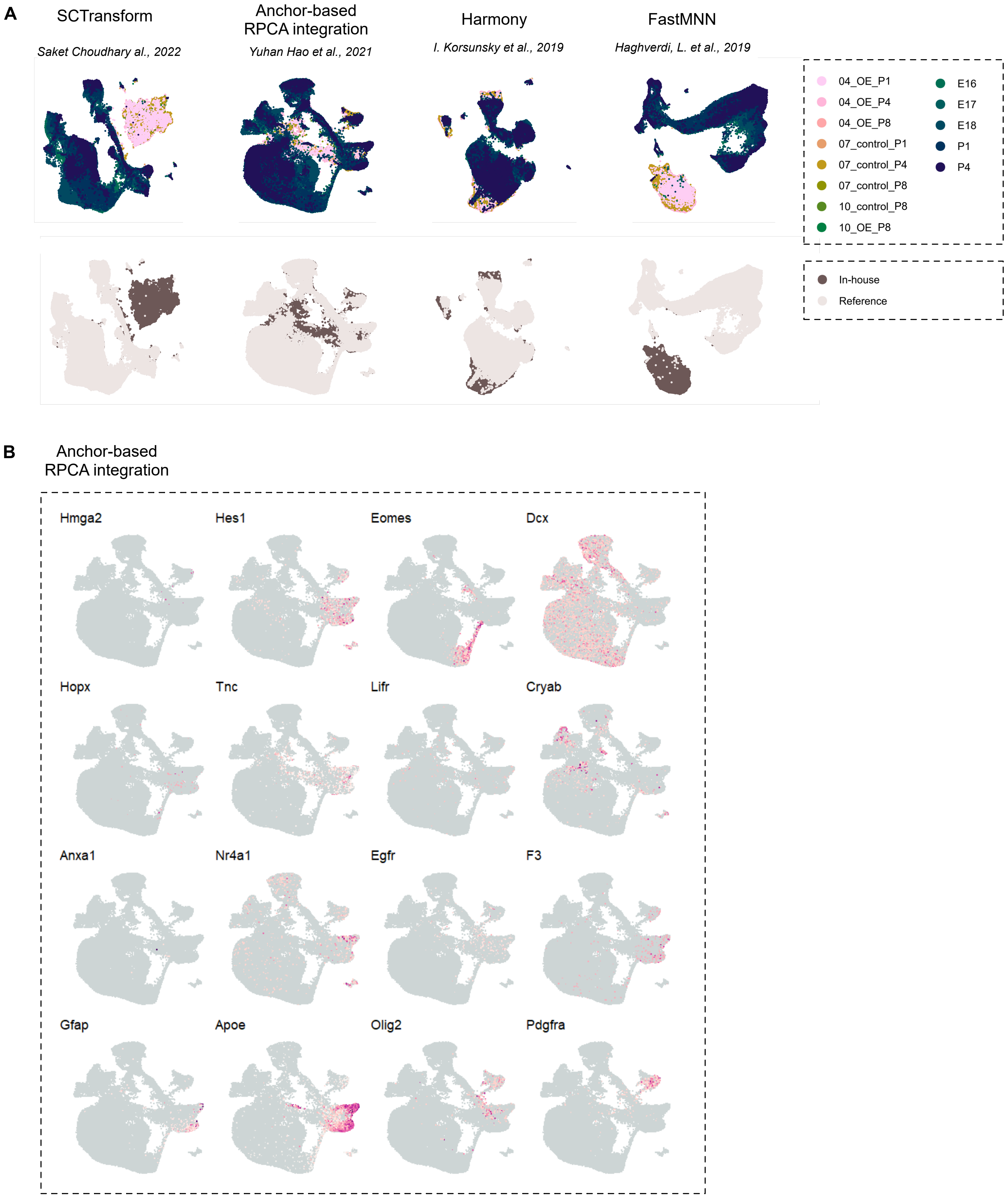

### FigS7

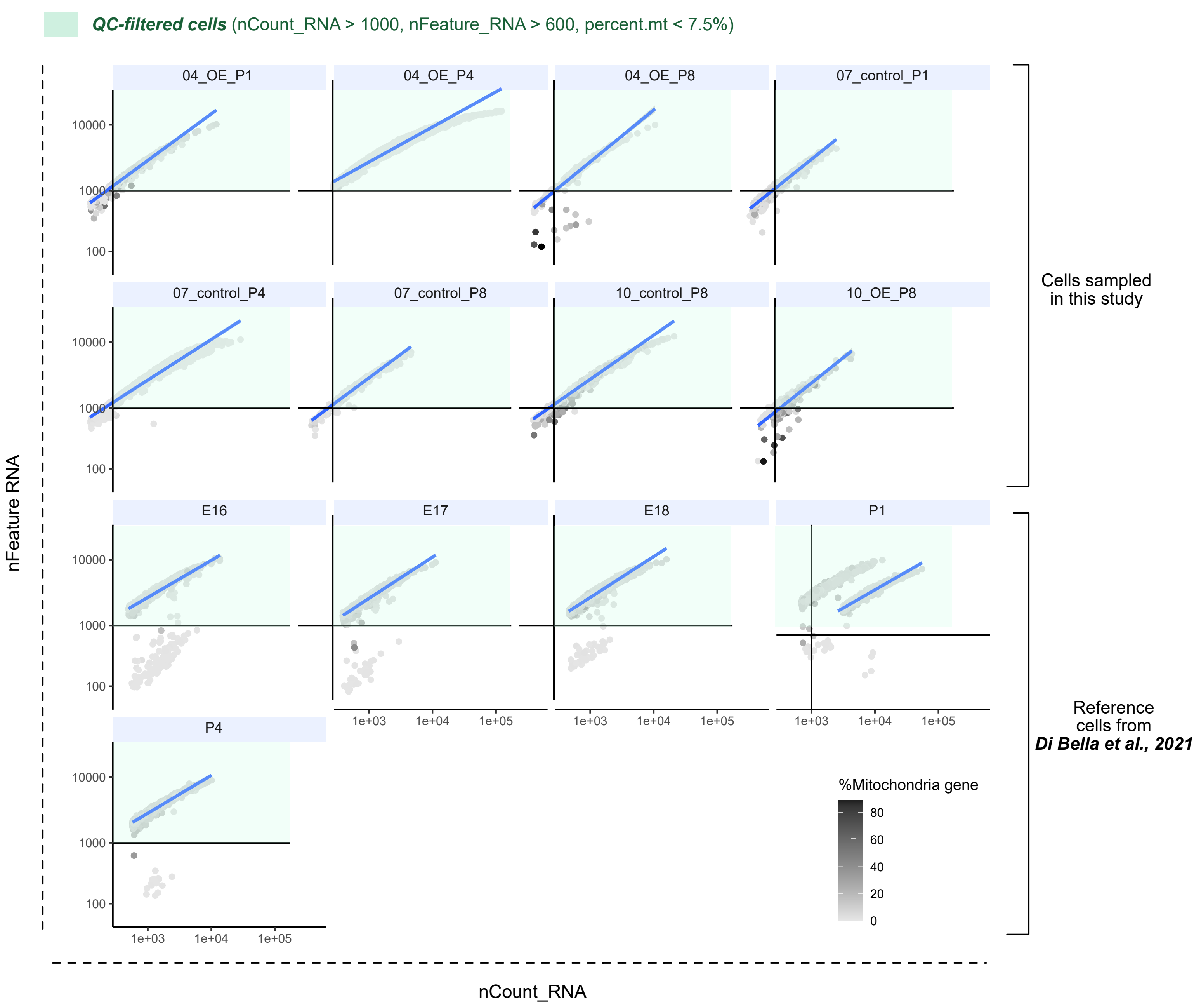
